# *IUCNN* - deep learning approaches to approximate species’ extinction risk

**DOI:** 10.1101/2021.06.17.448832

**Authors:** Alexander Zizka, Tobias Andermann, Daniele Silvestro

## Abstract

**Aim:** The global Red List (RL) from the International Union for the Conservation of Nature is the most comprehensive global quantification of extinction risk, and widely used in applied conservation as well as in biogeographic and ecological research. Yet, due to the time-consuming assessment process, the RL is biased taxonomically and geographically, which limits its application on large scales, in particular for understudied areas such as the tropics, or understudied taxa, such as most plants and invertebrates. Here we present *IUCNN*, an R-package implementing deep learning models to predict species RL status from publicly available geographic occurrence records (and other traits if available).

**Innovation:** We implement a user-friendly workflow to train and validate neural network models, and subsequently use them to predict species RL status. *IUCNN* contains functions to address specific issues related to the RL framework, including a regression-based approach to account for the ordinal nature of RL categories and class imbalance in the training data, a Bayesian approach for improved uncertainty quantification, and a target accuracy threshold approach that limits predictions to only those species whose RL status can be predicted with high confidence. Most analyses can be run with few lines of code, without prior knowledge of neural network models. We demonstrate the use of *IUCNN* on an empirical dataset of ∼14,000 orchid species, for which *IUCNN* models can predict extinction risk within minutes, while outperforming comparable methods.

**Main conclusions:** *IUCNN* harnesses innovative methodology to estimate the RL status of large numbers of species. By providing estimates of the number and identity of threatened species in custom geographic or taxonomic datasets, *IUCNN* enables large-scale analyses on the extinction risk of species so far not well represented on the official RL.

## Introduction

In face of the global biodiversity crisis, the disciplines of biogeography and (macro)ecology can provide an urgently needed perspective for global conservation (Brooks et al., 2006). Particularly promising contributions include the prioritization of species and areas for conservation and a mechanistic understanding of species extinction risk (Pollock et al., 2020; Rapacciuolo, 2019). Indeed, macroecological studies increasingly explore these topics, yet often limited to well studied regions or taxa, because available information on species extinction risk is scarce and geographically and taxonomically biased (Bachman et al., 2019; Donaldson et al., 2016).

The Red List of the International Union for the Conservation of Nature (RL, www.iucnredlist.org) is arguably the most influential scheme quantifying species extinction risk (Betts et al., 2020) and a prime example for the inclusion of biogeographic principles into conservation. The RL classifies species into five extinction risk categories: Least Concern (LC), Near Threatened (NT), Vulnerable (VU), Endangered (EN), and Critically Endangered (CR). Professional assessors or specialist groups comprised of volunteer scientists classify species into these categories (“red-listing”) based on at least one of five standardized criteria related to the reduction of population size (Criterion A), limited or shrinking geographic range (B), small population size and decline (C and D) or following a quantitative extinction risk assessment (E, IUCN, 2012; IUCN Standards and Petitions Subcommittee, 2017). Species that cannot be classified into any of these categories, because too little information is available, are considered Data Deficient (DD); species never considered in a red-listing process are termed Not Evaluated (NE).

Although the RL is designed for applied conservation, the standardized assessment across regions and taxa make it a valuable and widely-used resource for biogeographic and (macro)ecological research. For instance, RL extinction risk assessments have been used to relate traits, such as body mass, to species extinction risk (Boehm et al., 2016; Pincheira□Donoso et al., 2021; Richards et al., 2021; Rolland & Salamin, 2016), to quantify the effect of threats, such as agriculture, on species extinction risk (Polaina et al., 2018), to quantify links between species extinction risk and invasive species (Tingley et al., 2016; Walsh et al., 2012), to characterize the distribution of threatened species (Coll et al., 2015), to predict future biodiversity losses (Andermann et al., 2021; Monroe et al., 2019), and to understand the potential effects of extinction on large-scale diversity patterns (Oliveira et al., 2020; Smiley et al., 2020).

While the above mentioned examples illustrate the potential of the RL for biogeographic and macroecological research, the taxonomic and geographic biases of the RL prevent further integration, and limit research to well studied taxa or regions. Red-listing is time consuming due to the standardized process and the data requirements (assessment of a single species may take at least one day) which is why only a fraction of the global biodiversity has been evaluated with varying coverage across taxa. For instance, most known vertebrate species (68%) have been evaluated at least once, but the proportion of plants (7%), invertebrates (2%) and fungi & protists (<1%) is considerably lower (Bachman et al., 2019; IUCN, 2018; Lughadha et al., 2020). Furthermore, the proportion of evaluated species is higher in regions with experts and funding available (Bachman et al., 2019), and many of the existing assessments are, or will soon be, older than 10 years and thereby, outdated (Rondinini et al., 2014).

To speed up red-listing and to overcome said biases, a variety of methods have been developed to automate the red-listing process in recent years. These automation methods trade case-by-case evaluation for reproducibility, scope, and speed and they may process thousands of DD or NE species based on publicly available data within minutes (Zizka, Silvestro, et al., 2021). There are different flavors of automation methods, which differ in scope, underlying algorithms and data requirements. The most general approaches may infer indices required during the formal red-listing procedure, such as the Extent of Occurrence or the Area of Occupancy to support RL assessors (e.g., Bachman et al., 2011), provide preliminary RL assessments based on readily available data following IUCN criteria (e.g., Dauby et al., 2017), or predict species RL category based on species traits (e.g., González-del-Pliego et al., 2019; Pelletier et al., 2018).

While all of these automation approaches have important limitations (Lughadha et al., 2019; Rivers et al., 2011; Walker et al., 2020), they constitute useful tools for filling gaps in large datasets containing fractions of DD and NE species. Predictive approaches are particularly promising, since they are able to integrate different data types. For instance, predictions may be based on species geographic distribution, morphology and physiology as well as human disturbance, human use, and molecular data (Pelletier et al., 2018; Zizka, Silvestro, et al., 2021). Additionally, predictive approaches can benefit from the active development and ever-improving performance of novel machine learning methods. Among them, neural networks are a highly flexible family of models used to perform classification tasks or parameter estimation (LeCun et al., 2015). Deep neural networks have been shown to be able to approximate virtually any function, thus providing one of the most general and powerful available frameworks for predictions (Goodfellow et al., 2016). While some of the automated assessment methods are implemented in accessible software, most are not. To our knowledge, no application to use neural networks for RL prediction exists. More generally, the few existing attempts to use machine learning for red-listing are documented in scripts from the supplementary material of research studies difficult to access for the broader community and to adapt to different taxonomic and geographic scopes.

Starting from a deep learning method we recently applied to predict the RL status of orchid species globally (Zizka, Silvestro, et al., 2021), we here present *IUCNN*, an R-package implementing multiple neural network algorithms to predict species RL status in an accessible, user-friendly, and reproducible way.

## Methods

A typical workflow in *IUCNN* contains three major steps: data preparation, model training and testing, and prediction and visualization, which can be run as a pipeline (Fig. 1) with few lines of code (Fig. 2).

**Figure 1.**
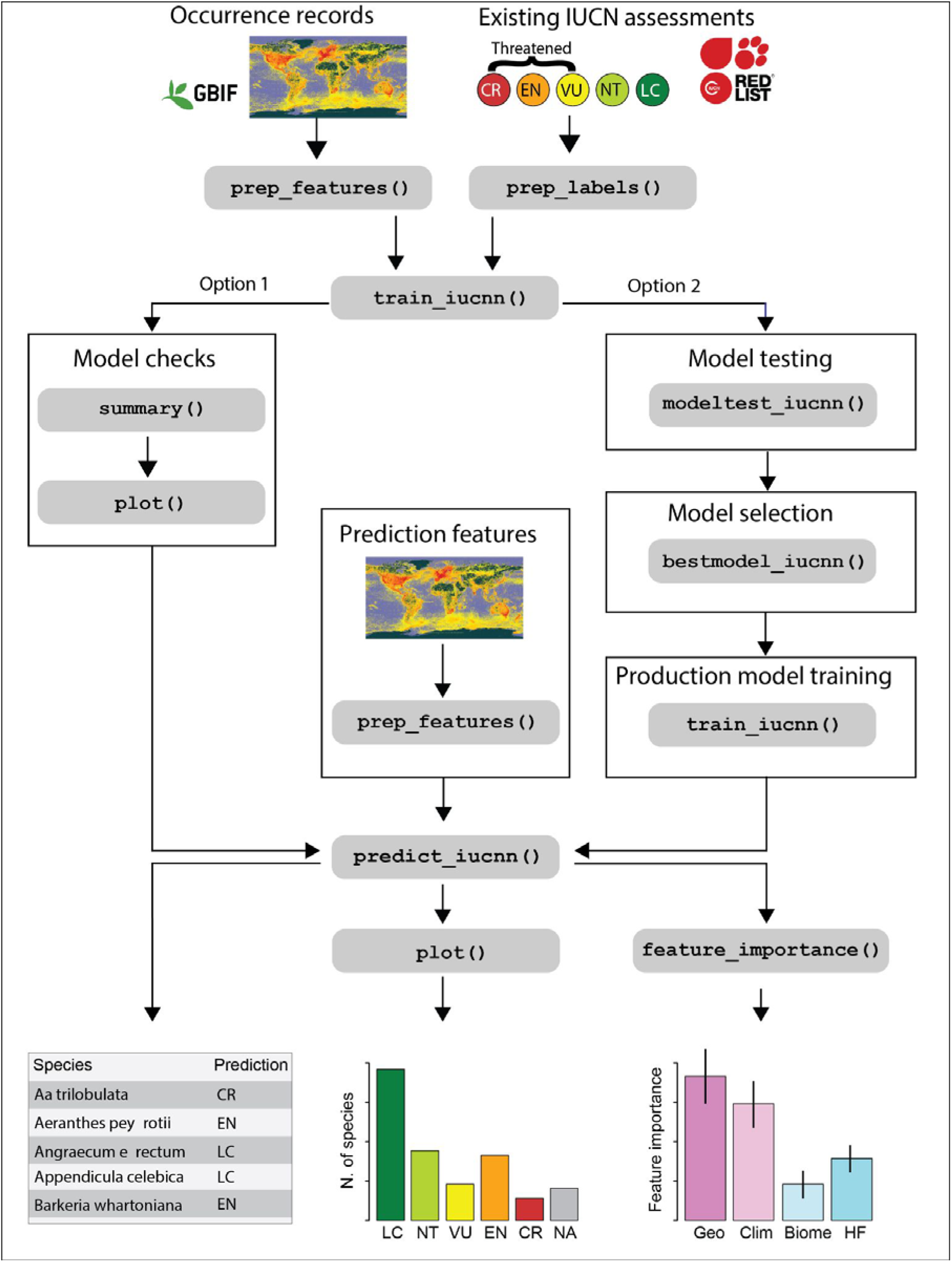
Schematic diagram of an *IUCNN* workflow. The relevant *IUCNN* functions are shown in the grey boxes.

**Figure 2.**
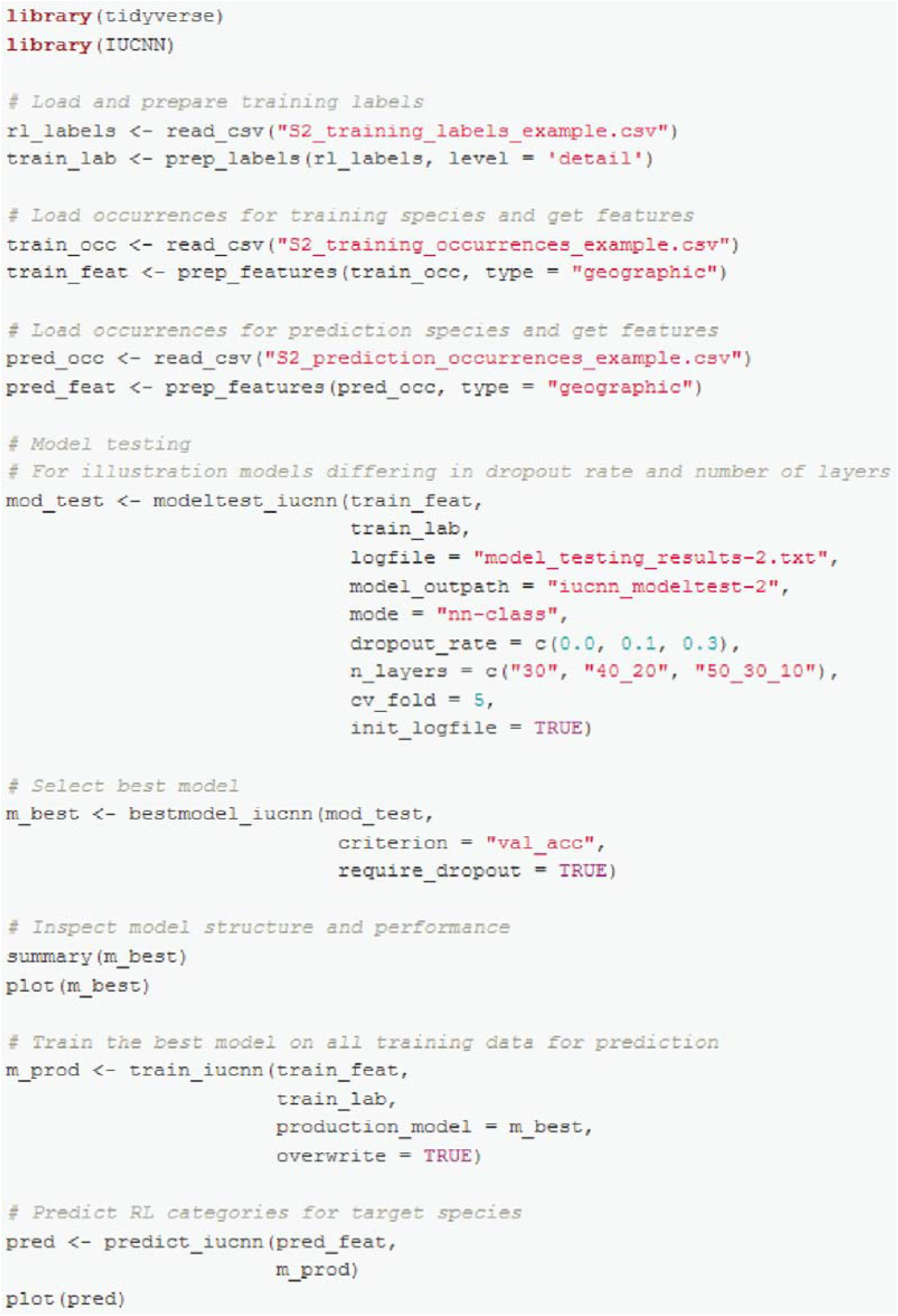
Example code on how to train and test a neural network using *IUCNN* and use it to predict the RL status of 13,207 orchid species, including model testing and selection (“Option 2” in Figure 1). The example data used in the displayed code are available in Supplementary material S3 and reduced example datasets are provided as part of the package.

The *IUCNN* package implements different types of neural network models and a set of functions for a reproducible and accessible workflow. Specifically, *IUCNN* implements well documented functions to:

1. calculate standardized geographic, climatic, and ecological features that can be used for RL extinction risk predictions from user-provided species geographic occurrence records
2. train deep learning models
3. predict species RL extinction risk category and quantify the uncertainty around the predictions
4. evaluate the importance of the different input features in determining the predictions

### Label and feature preparation

*IUCNN* models use features derived from species traits and are trained on species already assessed in the RL. Features may derive from any type of trait but in the simplest case publicly available species occurrence records suffice. Thus, the only input data needed are a set of species with existing RL assessments (the training labels), geo-referenced occurrence records for these species (from which the training features are extracted), and geo- referenced occurrence records for the species whose RL status is to be predicted.

*IUCNN* contains two functions to facilitate label and feature preparation. The `prep_label` function prepares standardized labels (i.e. the five IUCN categories, or a binary classification: Possibly threatened/Not threatened) in the format needed for model training. In the optimal case, models will be trained on existing RL assessments from evolutionarily related or ecologically similar species (e.g. all members of the orchid family worldwide, to predict extinction risk of orchid species) or related species in the same geographic region (e.g. vascular plants in South America, to predict the extinction risk of South American orchids). All existing RL assessments are publicly available online (https://www.iucnredlist.org/) and may be obtained in R using the rredlist package (Chamberlain, 2020). In general, the more species are in the training set, the better the model performance, and we recommend at least several hundred species for training, with a balanced representation of different RL categories as far as possible.

While IUCNN models can be trained on any features that might be informative of species RL category, for many species only geographic occurrence data are readily available. Thus, we implemented a `prep_features` function to extract and prepare a set of default features from user-provided geo-referenced occurrence records of species. The function automatically downloads publicly available environmental data and matches these to the user-provided occurrences. These default features capture several geographic characteristics of each species range. Additionally, for terrestrial taxa, the default features include the intensity of human impact within the species range (Venter et al., 2016, obtained from https://wcshumanfootprint.org) the climatic conditions across a species range (Fick & Hijmans, 2017, obtained from https://www.worldclim.org via the raster package), and species occurrences in different biomes (Olson et al., 2001, obtained from http://assets.worldwildlife.org). We provide a detailed list of all features in Table S1 in Supplementary material S1. The download of environmental data, as well as the subsequent data extraction, summary, and standardization are handled by the `prep_features` function, but may be customized by the user. In case no custom dataset of geo-referenced species occurrences is available, these may be obtained from public databases, for instance the Global Biodiversity Information Facility (GBIF). Records from GBIF can be downloaded from www.gbif.org or obtained via the rgbif package in R (Chamberlain & Boettiger, 2017). If data from an online database are used, taxonomic scrubbing (Cayuela et al., 2012; Freiberg et al., 2020), quantification of sampling bias (Zizka et al., 2021) and geographic cleaning (Zizka et al., 2019) are advisable (Maldonado et al., 2015; Zizka, Carvalho, et al., 2020).

Users may also provide a dataset of custom features. Features may be continuous or discrete and may represent any trait considered relevant for approximating the conservation status in the region and taxon of interest, such as phenotypic data or population dynamics. Features should be rescaled so that their numeric values range in similar orders of magnitude to facilitate model convergence, and missing data should be coded with distinct values or imputed, for instance using the missForest R-package (Stekhoven & Bühlmann, 2012). When using custom features it is important that the same features are provided in the training and prediction datasets, and that they are rescaled consistently.

### Model training and testing

We implemented three models in *IUCNN*, all of which are based on fully connected neural networks (NNs), with user-defined architecture and hyper-parameters, which we term: *nn-class, bnn-class*, and *nn-reg*. The *nn-class* model is a classifier built and optimized through the Tensorflow module (v2 or greater; Abadi et al., 2015), the *bnn-class* model is a Bayesian implementation of an NN classifier using the npBNN module (Silvestro & Andermann, 2020), and the *nn-reg* model predicts the conservation status of species as a regression task, also using Tensorflow. The model training, although based on different libraries, is packaged in a consistent and user-friendly framework within *IUCNN*, and the user is not required to know how to use such libraries running under the hood.

In all models, the input layer consists of a set of quantitative and categorical features computed for each species (see above). The input features are mapped onto the output layer through one or more hidden layers; the number of hidden layers, the number of nodes per layer, and the activation functions can be adjusted by the user. The *nn-class* and *bnn-class* models are classifiers and use a SoftMax activation function in the output layer to obtain a vector of probabilities, with one value for each class, e.g. the five RL categories. The prediction under these models is determined by the class that received the largest probability value. In the *nn-class* model, the probabilities are then used to compute the cross-entropy loss, which is averaged across all training instances and minimized during the optimization. In the *bnn-class* model the output class probabilities are used as the parameters of a categorical probability mass function to compute the likelihood of the training data. The output layer of the *nn-reg* model consists of a single value, which may be taken as is, or transformed through a sigmoid or tanh activation function (depending on user-defined settings for the output activation function). The output value is then compared with the original or rescaled RL status during model optimization as mean squared error (MSE). *IUCNN* then transforms the predicted value into a categorical prediction by rounding the output to the closest class.

The three models are conceptually different, with individual advantages and limitations. The *nn-class* and *bnn-class* models do not explicitly incorporate the ordinal nature of the RL categories, treating them instead as categorical. The potential disadvantage of this approach is that the error is not weighted by the degree of disagreement between the truth and the prediction. For instance, assessing a NT species as LC or CR will be considered as equally wrong, even though NT is much closer to LC than to CR. To overcome this limitation, the classes are handled differently in the *nn-reg* model, which optimizes the output as a regression, where the distance (error) between NT and LC is much smaller than between CR and LC. Thus, while the accuracies are expected to be similar among the three models, the magnitude of the error in erroneously classified instances is expected to be smaller in the *nn-reg* model.

The *bnn-class* model is a Bayesian implementation of the *nn-class* model, with the advantage of producing a posterior sample of predictions for each instance instead of a point estimate. From these samples we compute posterior probabilities associated with each class, thus providing a direct estimation of the uncertainty around the prediction. As demonstrated in the empirical example, the posterior probabilities can be used to determine a threshold above which instances are expected to be classified with an accuracy matching or exceeding a user-defined target. The advantage of the BNN implementation comes at the cost of a time consuming optimization. To also quantify the uncertainties of the predictions made by the *nn-class* and *nn-reg* models, we implemented the Monte Carlo dropout method (Gal & Ghahramani, 2016). While the resulting dropout probabilities are not posterior probabilities, they can be similarly applied and interpreted as a measure of prediction uncertainty and allow users to determine for which species a prediction can be made with a defined level of confidence.

The *nn-class* and *nn-reg* models are optimized using the Adam gradient descent optimization (Kingma & Ba, 2017) as implemented in Tensorflow. The *bnn-class* model is trained through a Markov Chain Monte Carlo (MCMC) algorithm, sampling the weights from their posterior distribution. By default, standard normal priors are applied to the weight parameters. The user can define a number of parameters controlling the training process (e.g., the number of epochs, stopping criteria for *nn-class/nn-reg*, and the number of MCMC iterations and the sampling frequency for *bnn-class*) through the function `train_iucnn`.

The performance of trained *IUCNN* models can be evaluated using the `summary` function, which calculates summary statistics for the model on unseen data including an overall prediction accuracy and a confusion matrix. Additionally the `plot` function can be used to plot the loss of the trained *IUCNN* model throughout the training epochs (*nn-class* and *nn-reg*) or the posterior samples (*bnn-class*), which helps to evaluate whether or not the model has converged. To evaluate a range of different model settings we implemented the `modeltest_iucnn` function, which allows the user to choose among different NN architectures and hyperparameter configurations (e.g. number of layers, nodes per layer, activation functions) using a cross-validation approach. Finally, users can evaluate how much the chosen model relies on the different types of features, using the `feature_importance` function. The implemented process of permutation feature importance (Breiman, 2001), evaluates the loss of prediction accuracy when the signal in individual features is muted by randomly shuffling the feature values among instances. The resulting feature importance values for each feature or block of features can be plotted with the `plot` function. This can help users to decide which features are most important and should be included in a model to obtain the best prediction accuracy (Fig. 1, Table 1). We provide a tutorial on how to train and test models in *IUCNN* as well as a vignette that comes with the R-package.

**Table 1.**
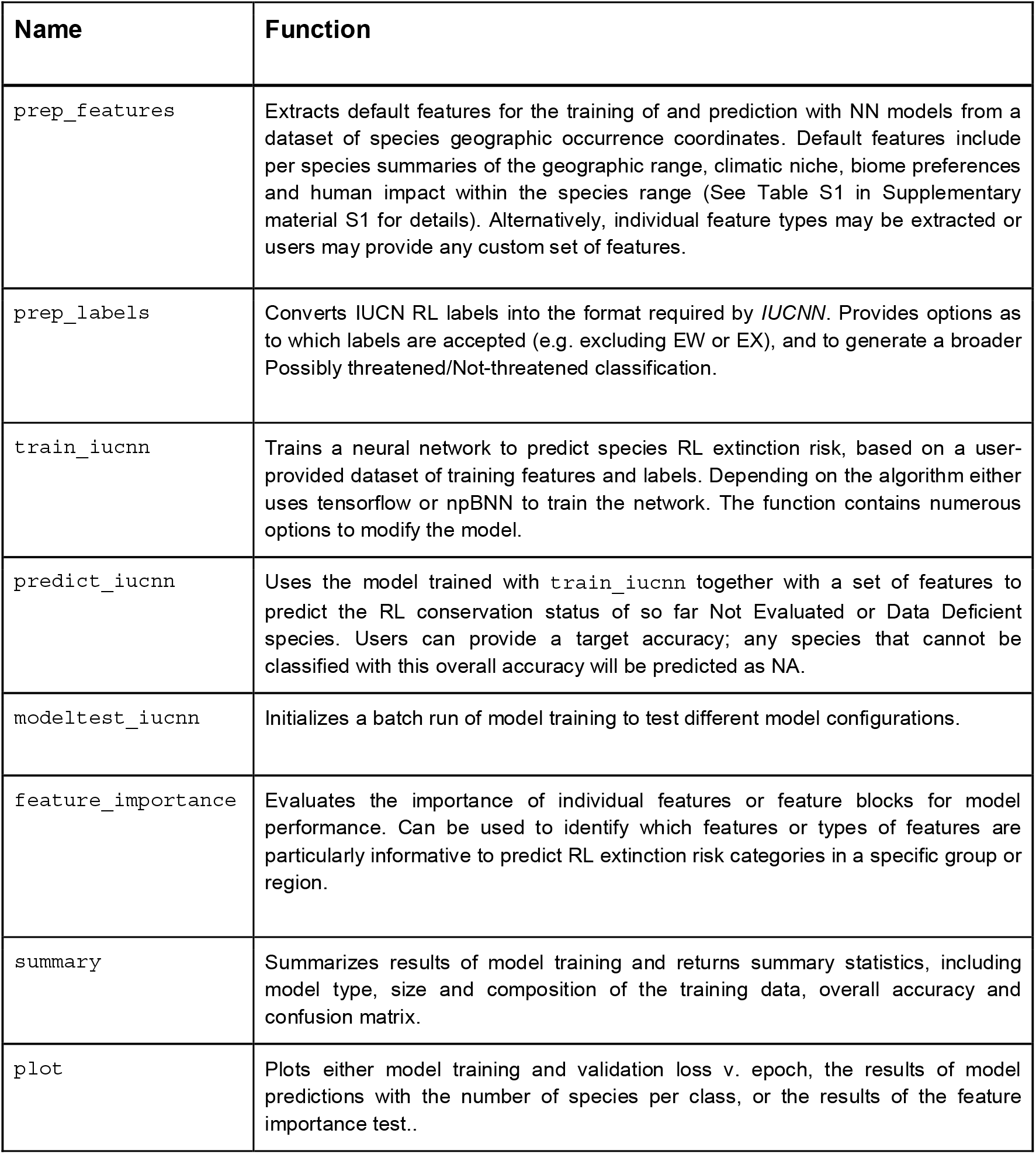
The main functions of the *IUCNN* package. An example pipeline on how to use these functions is shown in Figure 1 and the options to customize analyses are detailed in the vignette distributed with the package.

### Prediction and visualization

Provided with the trained model and the features for the target species, the `predict_iucnn` function predicts RL categories for these species. Labels may be returned as a vector of labels or optionally, in the case of an *nn-reg* analysis, as the raw regressed values. In case a target accuracy was set, all species for which the RL category cannot be predicted at the desired accuracy are labeled as NA. Using the `plot` function, users can plot a histogram of the numbers of predicted species per category.

### Implementation

*IUCNN* is implemented as an R-package (R Core Team, 2021) with integrated Python code (using the Tensorflow and npBNN modules). All user-level functions are accessible via R, since this language is widely used in ecological and conservation research and no Python knowledge is required to use *IUCNN*. We developed *IUCNN* using the ‘usethis’ package (Wickham & Bryan, 2020) following Wickham and Bryan (2021).

*IUCNN* depends on the R-packages dplyr (Wickham, François, et al., 2020), magrittr (Bache & Wickham, 2014), readr (Wickham & Hester, 2020), tidyr (Wickham, 2020), and tidyselect (Henry & Wickham, 2020) for data wrangling, raster (Hijmans, 2018), rCAT (Moat, 2017), sf (Pebesma, 2018), and stats for feature preparation, graphics and grDevices for visualization, and reticulate (Ushey et al., 2020) for integrating R and Python. Furthermore, *IUCNN* suggests checkmate (Lang, 2017), covr (Hester, 2020), spelling (Ooms & Hester, 2020), testthat (Wickham, 2011) to secure code functionality and knitr (Xie, 2020) and rmarkdown (Allaire et al., 2020) for documentation.

All software needed to run *IUCNN* can be installed with a few lines of code. The current version of *IUCNN* can be installed from GitHub (https://github.com/azizka/IUCNN), from within R using the devtools (Wickham, Hester, et al., 2020) package. Since the neural networks are trained and used in Python, Python also needs to be installed, including the tensorflow module. This can be done from within R using the reticulate package (Ushey et al., 2020). See the readme file on IUCNN’s GitHub page or the vignette provided with the package (Supplementary material S2) for the necessary code for installation as well as an empirical data tutorial.

## Results

We use an empirical dataset on the global distribution and extinction risk of orchid species (Orchidaceae) to demonstrate the use of *IUCNN*. The dataset contains 14,093 orchid species from across the globe, 886 of them with existing RL assessments (the training data). The specific dataset is described in detail in Zizka, Silvestro et al. (2021), and the occurrence records are originally obtained from GBIF (Global Biodiversity Information Facility (www.gbif.org), 2019). The dataset has already been processed with the ConR (Dauby et al., 2017), rCAT (Moat, 2017), and SPGC (Schmidt et al., 2017) automation methods in Zizka, Silvestro et al. (2021), which we use for comparison. These three methods are comparable to *IUCNN* in that they are also solely based on species geographic occurrence records and are implemented in accessible R-packages, but differ in that they do not predict extinction risk based on traits, but calculate RL indices (Extent of Occurrence and Area of Occupancy) relevant for red-listing under Criterion B, and may be interpreted as preliminary assessment.

To illustrate the use and flexibility of *IUCNN*, we train and test *nn-class, nn-reg*, and *bnn-class* models based on different sets of features, with different model structures and different levels of prediction detail (5 RL categories and Possibly threatened v. Not threatened species, respectively). We tested *nn-class and nn-reg* models based on geographic features, geographic plus human footprint features and all default features. We compared different *nn-class* models with 1, 2, and 3 hidden layers and dropout rates set to 0, 0.1, and 0.3 (detailed settings are available in the accompanying vignette distributed with the package and Supplementary material S2). For *nn_reg* models we tested the same specifications, and included different output layer activation functions. We tested all combinations of these settings. Within each model type and output classification, we chose the best model based on the cross-validation accuracy. Due to the longer convergence time we ran the *bnn-class* models assuming a single configuration with one hidden layer with 20 nodes.

The overall accuracy of models was similar across the tested subsets of input features. In general, models based solely on geographic features performed best (in 2 out of 4 cases, more often than any other feature combination, Fig. S1 in Supplementary material S1), and we therefore used these models for further analyses.

The accuracy of any *IUCNN* model was higher than the accuracy of the available automation methods based on Criterion B (Fig. 3A). For all *IUCNN* models, the overall accuracy was higher at the Possibly threatened vs. Not threatened level (nn-class: 0.81, nn-reg: 0.80, bnn-class: 0.80; Fig. 3A) than at the detailed level with all five RL categories (nn-class: 0.60, nn-reg: 0.54, bnn-class: 0.60; Fig 3A). While the overall accuracy was slightly higher for the *nn-class and bnn-class* model, the *nn-reg* model performed better in including intermediate categories, in particular NT and VU (Fig. 3B). Given that most species fall into the LC category, this advantage of the nn-reg model was not reflected in the overall prediction accuracy. When using target accuracy thresholds, the proportion of species evaluated decreased with increasing target accuracy for all model types and detail levels (Fig. 4). With increasing target accuracy, species of intermediate categories are subsequently removed and species that can be classified at higher accuracies mostly belong to the extreme LC and CR categories (Fig. S2 in Supplementary material S1). In the case of the best *nn-class* model, a target accuracy of 80% retains 3,310 species (out of 13,207), 3004 of them as LC, 104 as EN and 202 as CR.

**Figure 3.**
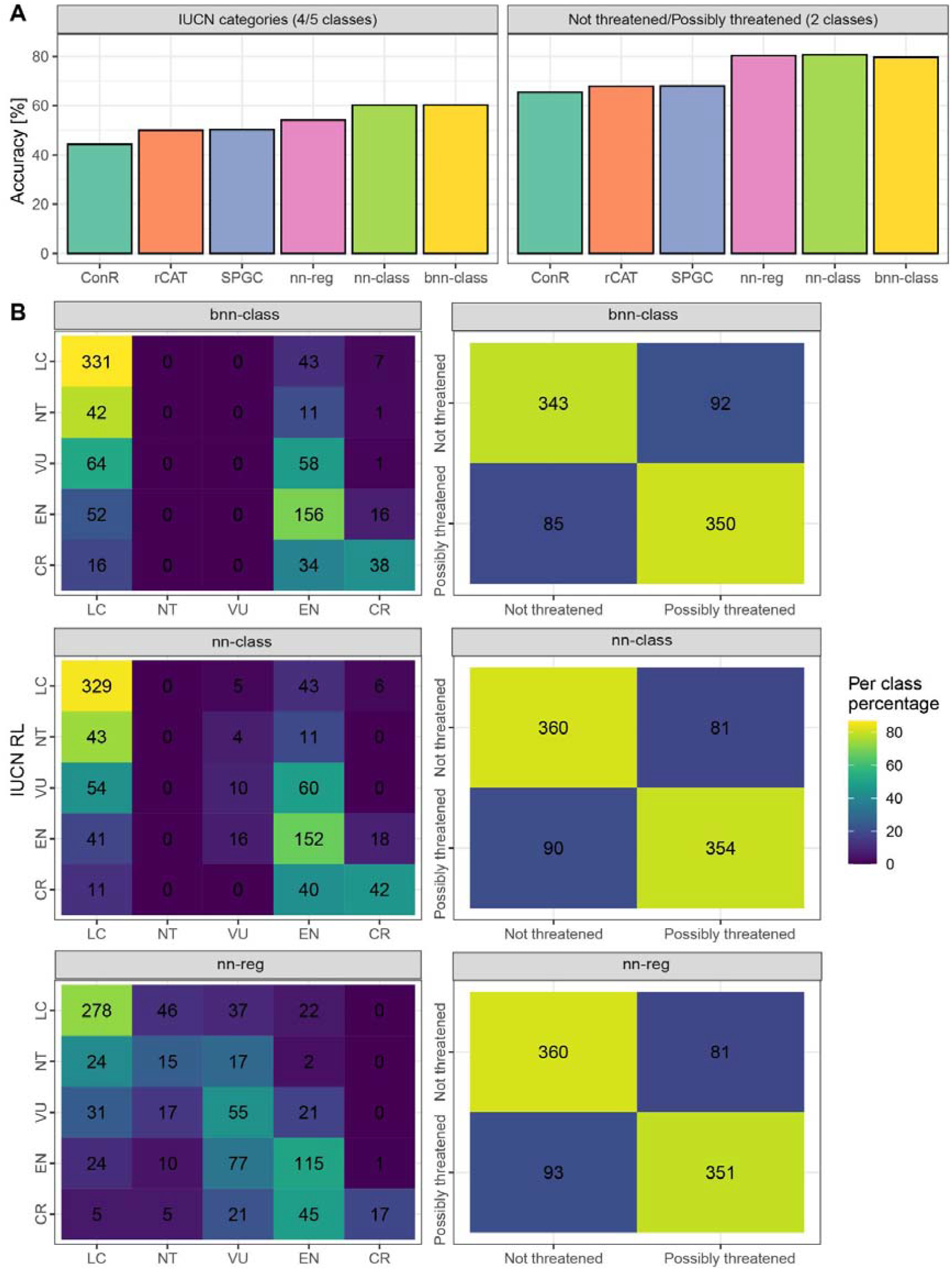
The performance of different *IUCNN* models in predicting the extinction risk of 13,207 orchid species trained on 886 species with existing RL assessments at www.iucnredlist.org. **A)** The overall accuracy of different automated methods to estimate the extinction risk of species on the level of RL categories or the broader Possibly threatened/Not threatened level. ConR, rCat and SPGC are other automation methods for comparison. **B)** Confusion matrices of the three *IUCNN* models at the detailed RL category and the broad Possibly threatened/Not threatened level.

**Figure 4.**
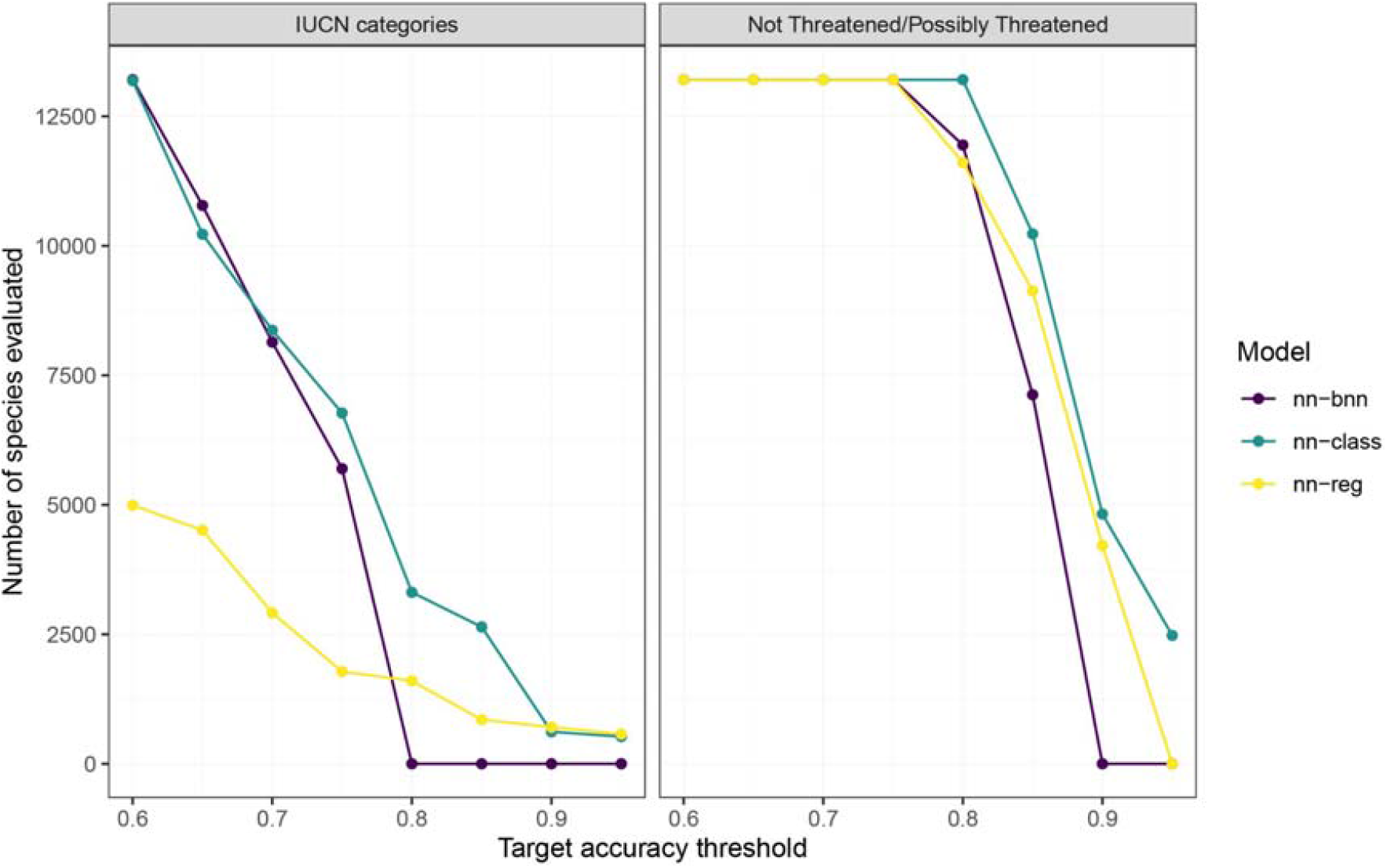
The effect of different user-defined targets for overall model accuracy on the number of species assessed. Based on the example dataset of 13,207 orchid species with no official assessment available. The number of assessed species decreases with increasing accuracy, but in this example the *nn-class* model can classify more than 3,000 species at 80% accuracy.

## Discussion

We presented *IUCNN*, a user-friendly R-package to use neural networks for the prediction of species RL assessments. *IUCNN* is flexible with regard to input data and may be used solely based on publicly available information. Our empirical example using more than 13,000 orchid species demonstrates that the models implemented in *IUCNN* can be run with few lines of code, and can outcompete other automated methods in terms of overall accuracy.

Predictive models, such as the neural networks implemented in *IUCNN*, provide approximations of species RL assessments, which can be used in biogeographic and (macro)ecological research. Specifically, these approximations are suitable to broaden the taxonomic and geographic scope of synthetic research into understanding the distribution and mechanisms of threat and extinction risk. When extinction risk is used, for instance, to understand the geographic distribution of threatened species or to quantify the impact of species traits on extinction risk, predictive approaches can overcome taxonomic and geographic biases, and enable such analyses for organisms that are poorly represented in the current RL. Furthermore, predictive approaches can support ecological, biogeographic and taxonomic case studies that can benefit from information on species extinction risk. For example, added information on the estimated number and identity of threatened species in a study taxon or region may increase the conservation relevance of research, help to set research priorities or even guide ethical decisions on sampling strategies (for instance by switching to less invasive sampling techniques for Possibly threatened species).

Theoretically, predictive models as those implemented in *IUCNN*, may be one option to speed up the official red-listing process and overcome geographic and taxonomic biases by feeding into the RL directly. However, the uncertainty related to model predictions, common misapplication of official RL criteria in predictive methods, the conceptual inaccessibility of many predictive models for non-specialists, and their circumvention of the formal IUCN red-listing documentation and criteria make an integration unlikely (e.g., Akçakaya et al., 2006; Walker et al., 2020). A specific case where predictive models may indeed support applied red-listing is the funneling of expert time to Possibly threatened species. For instance, predictive approaches may be used as a first step to identify species that are most likely to be considered LC by a formal RL assessment and remove them from further inspection, thereby reducing the pool of species requiring expert attention and focus RL expert time to species that are likely to be classified at high risk (Bachman et al., 2020).

We designed *IUCNN* for user-friendly access. In addition, we included multiple features to facilitate the use cases of biogeographical and (macro)ecological research and to address common issues arising with the use of RL data prediction of extinction risk. Five specific strengths of *IUCNN* are:

### Easy model evaluation and model testing

Careful model evaluation and testing is crucial for all statistical models. *IUCNN* provides options for model evaluation, testing and customization. When predicting extinction risk and training on RL data, high accuracies are crucial (since errors may result in a misprioritization of conservation resources) and uncertainty must be clearly visible. The implementation of the *bnn-class* Bayesian neural network allows for a probabilistic quantification of uncertainty, and the implementation of a target accuracy allows limiting prediction to high levels of accuracy. To address the limited amount of training data, the implementation of cross-validation for model testing and the option to subsequently train a model on all available data ensures to use as much of the existing training data as possible.

### High accessibility

*IUCNN* is implemented in R, a widely-used programming language in ecology, and an individual analysis may be run with few lines of code in a straightforward workflow (Fig. 1, Fig. 2). Although a Python installation is required, the installation process can be navigated from within R. *IUCNN* contains convenient functions to prepare standardized features in the required input format suggesting a set of default features that have proven relevant in empirical analyses (`prep_features`) and easy options to customize them via separate functions (`ft_biom, ft_clim, ft_foot, ft_geo`). Model testing and evaluation are concentrated in just three functions (`modeltest_iucnn, bestmodel_iucnn,` and `feature_importance`) and the results can easily be summarized and visualized via the standard `plot` and `summary` methods. All functions are documented at the standards of the Comprehensive R Archive Network and the *IUCNN* workflow as well as the different options for customization are detailed in the accompanying vignette distributed with the package. Furthermore, the majority of *IUCNN* analyses (including several thousands species and hundreds of features) can be run on a standard laptop.

### High flexibility

*IUCNN* may predict species extinction risk based on any trait deemed relevant by the users, as long as they can be summarized at the species level. While we provide custom features that can be derived from data publicly available for many taxa, users can easily provide data describing for instance the physiology, taxonomy, morphology, ecology of species. This flexibility can avoid circularities when for example using *IUCNN* predictions to relate extinction risk to specific traits. Furthermore, *IUCNN* may use any user-desired output labels. These may be the default five IUCN categories (LC, NT, VU, EN, CR) but may also be any user-defined classes, such as the binary classification demonstrated in the empirical example, or for instance classification schemes of regional Red Lists, which often are different.

### Improved accuracy and future prospects

Neural networks can outcompete comparable methods to identify possibly threatened species and to classify species into RL categories, as demonstrated by the empirical analysis (Fig. 3A). Neural networks and deep learning are an active field of development, with multiple lines of development on the horizon. Together with the growing amount of RL data that can be used for training and the increase of remote sensing technologies for feature preparation, we anticipate an increase in application cases and accuracy of *IUCNN* models.

While we are confident that these specific adaptations of *IUCNN* make it a useful tool for research, we emphasize that trait-based predictions of species extinction risk are only approximations and need to be interpreted with caution (Walker et al., 2020). As for any statistical model, the quality of *IUCNN* predictions will depend on the quality of the input data and model fit. In the specific RL framework, the structure of the RL likely limits the maximum accuracy that can be achieved in predicting species extinction risk based on traits. The RL is a collaborative effort of many different specialist groups and professional assessors generating the assessments for individual taxonomic groups. Although assessments are based on a standardized framework, different assessors may interpret criteria and categories differently, especially the less clearly defined NT and VU categories. This effect is exacerbated by the differences in data availability among taxa and the resulting varying reliability of individual assessments. Furthermore, species may be listed on the RL based on different criteria, for instance either geographic range (Criterion B) or population trends (Criterion A), or both and assessors include expert knowledge and evaluation into the assessment. Hence, any set of input features may only incompletely capture the red-listing process and therefore misclassify species.

A specific caveat of predicting RL categories is their imbalanced distribution. In our empirical example, the models accurately identified species in the extreme categories (LC and CR), but performed less well with intermediate categories—the majority of misclassifications were related to NT and VU categories (Fig. 3B). This is likely related to these categories being relatively rare on the RL and thus underrepresented in the training data, which may bias predictions to reproduce category frequencies observed in the input data. Indeed, the models used for the empirical data were biased towards class label frequencies in the training data, especially if the frequencies were very different between training and test sets (Fig. S3 in Supplementary material S1). The implications of class imbalance for model application will depend on the expected similarity in class frequencies among species already evaluated on the RL (the training data) and DD and NE species (the target species). Conceptually, the expected difference in category frequencies remains unclear, because different factors affect the class frequencies of RL categories in a given set of species. On the one hand, by design, most species in any dataset will be LC, often also among the DD and NE species (Butchart & Bird, 2010). On the other hand, compared to species on the RL, datasets of DD and NE species are likely to comprise a higher proportion of rare species, which may be more threatened (Parsons, 2016). Yet, DD and NE species will more often occur in regions difficult to access for IUCN assessors (Bland et al., 2017) and might also be subject to reduced human pressure in general, and therefore have a reduced extinction risk. We acknowledge these conceptual and practical limitations for RL status predictions in general and have therefore included a battery of options to address these issues in *IUCNN*. These include the *nn-reg* models to account for the ordinal nature of the RL categories, the flexible detail levels in the prediction and the target accuracy threshold to reduce uncertainty.

In conclusion, *IUCNN* is a user-friendly implementation of deep learning methods to approximate RL extinction risk assessments for species that are so far Data Deficient or Not Evaluated on the RL. *IUCNN* provides new tools to estimate species extinction risk and makes these innovative methods available to a larger community and facilitates further integration of biogeography, (macro)ecology and conservation research.

## Supporting information

IUCNN Tutorial

Data for empirical analyses

## Acknowledgements

We thank the IUCN and all Red List assessors as well as the data collectors and data providers to GBIF for their effort. We are thankful to Steven Bachman and Barnaby Walker for helpful discussion during the development of IUCNN. AZ was funded by iDiv via the German Research Foundation (DFG FZT 118), specifically through sDiv, the Synthesis Centre of iDiv. T.A. and D.S. received funding from the Swedish Research Council (VR: 2019-04739). D.S. received funding from the Swiss National Science Foundation (PCEFP3_187012; FN-1749).

## Data availability statment

A stable version of *IUCNN* will be available as a release from https://github.com/azizka/IUCNN under a LGPL-2.1 license (v1.0.0), the developmental version and development history are freely available from GitHub under the same license. Example data and a detailed tutorial are available with the package.

## Supporting Information

Supplementary material S1 - Supplementary Figures and Tables

Supplementary material S2 - The vignette “Approximate_IUCN_Red_List_assessments_with_IUCNN” for *IUCNN* v1.0.0

Supplementary material S3 - Supplementary data to run the example code in Figure 2

## References

Abadi, M., Agarwal, A., Barham, P., Brevdo, E., Chen, Z., Citro, C., Corrado, G. S., Davis, A., Dean, J., Devin, M., Ghemawat, S., Goodfellow, I., Harp, A., Irving, G., Isard, M., Jia, Y., Jozefowicz, R., Kaiser, L., Kudlur, M., … Zheng, X. (2015). TensorFlow: Large-scale machine learning on heterogeneous systems. https://www.tensorflow.org/

Akçakaya, H. R., Butchart, S. H. M., Mace, G. M., Stuart, S. N., & Hilton□Taylor, C. (2006). Use and misuse of the IUCN Red List Criteria in projecting climate change impacts on biodiversity. Global Change Biology, 12(11), 2037–2043. https://doi.org/10.1111/j.1365-2486.2006.01253.x

Allaire, J. J., Xie, Y., McPherson, J., Luraschi, J., Ushey, K., Atkins, A., Wickham, H., Cheng, J., Chang, W., & Iannone, R. (2020). rmarkdown: Dynamic Documents for R. https://github.com/rstudio/rmarkdown

Andermann, T., Faurby, S., Cooke, R., Silvestro, D., & Antonelli, A. (2021). iucn_sim: A new program to simulate future extinctions based on IUCN threat status. Ecography, 44(2), 162–176. https://doi.org/10.1111/ecog.05110

Bache, S. M., & Wickham, H. (2014). magrittr: A forward-pipe operator for R. https://CRAN.R-project.org/package=magrittr

Bachman, S., Field, R., Reader, T., Raimondo, D., Donaldson, J., Schatz, G. E., & Lughadha, E. N. (2019). Progress, challenges and opportunities for Red Listing. Biological Conservation, 234, 45–55. https://doi.org/10.1016/j.biocon.2019.03.002

Bachman, S., Moat, J., Hill, A., Torre J. de la, & Scott, B. (2011). Supporting Red List threat assessments with GeoCAT: Geospatial conservation assessment tool. ZooKeys, 150, 117–126. https://doi.org/10.3897/zookeys.150.2109

Bachman, S., Walker, B., Barrios, S., Copeland, A., & Moat, J. (2020). Rapid Least Concern: Towards automating Red List assessments. Biodiversity Data Journal, 8, e47018. https://doi.org/10.3897/BDJ.8.e47018

Betts, J., Young, R. P., Hilton□Taylor, C., Hoffmann, M., Rodríguez, J. P., Stuart, S. N., & Milner□Gulland, E. J. (2020). A framework for evaluating the impact of the IUCN Red List of threatened species. Conservation Biology, 34(3), 632–643. https://doi.org/10.1111/cobi.13454

Bland, L. M., Bielby, J., Kearney, S., Orme, C. D. L., Watson, J. E. M., & Collen, B. (2017). Toward reassessing data-deficient species. Conservation Biology, 31(3), 531–539. https://doi.org/10.1111/cobi.12850

Boehm, M., Williams, R., Bramhall, H. R., McMillan, K. M., Davidson, A. D., Garcia, A., Bland, L. M., Bielby, J., & Collen, B. (2016). Correlates of extinction risk in squamate reptiles: The relative importance of biology, geography, threat and range size. Global Ecology and Biogeography, 25(4), 391–405. https://doi.org/10.1111/geb.12419

Breiman, L. (2001). Random Forests. Machine Learning, 45(1), 5–32. https://doi.org/10.1023/A:1010933404324

Brooks, T. M., Mittermeier, R. A., Fonseca, G. A. B. da, Gerlach, J., Hoffmann, M., Lamoreux, J. F., Mittermeier, C. G., Pilgrim, J. D., & Rodrigues, A. S. L. (2006). Global biodiversity conservation priorities. Science, 313(5783), 58–61. https://doi.org/10.1126/science.1127609

Butchart, S. H. M., & Bird, J. P. (2010). Data Deficient birds on the IUCN Red List: What don’t we know and why does it matter? Biological Conservation, 143(1), 239–247. https://doi.org/10.1016/j.biocon.2009.10.008

Cayuela, L., Cerda, Í. G. la, Albuquerque, F. S., & Golicher, D. J. (2012). taxonstand: An R package for species names standardisation in vegetation databases. Methods in Ecology and Evolution, 3(6), 1078–1083. https://doi.org/10.1111/j.2041-210X.2012.00232.x

Chamberlain, S. (2020). rredlist: “IUCN” Red List Client. https://CRAN.R-project.org/package=rredlist

Chamberlain, S., & Boettiger, C. (2017). R Python, and Ruby clients for GBIF species occurrence data. PeerJ PrePrints. https://doi.org/10.7287/peerj.preprints.3304v1

Coll, M., Steenbeek, J., Lasram, F. B. R., Mouillot, D., & Cury, P. (2015). “Low-hanging fruit” for conservation of marine vertebrate species at risk in the Mediterranean Sea. Global Ecology and Biogeography, 24(2), 226–239. https://doi.org/10.1111/geb.12250

Dauby, G., Stévart, T., Droissart, V., Cosiaux, A., Deblauwe, V., Simo□Droissart, M., Sosef, M. S. M., Lowry, P. P., Schatz, G. E., Gereau, R. E., & Couvreur, T. L. P. (2017). ConR: An R package to assist large-scale multispecies preliminary conservation assessments using distribution data. Ecology and Evolution, 7(24), 11292–11303. https://doi.org/10.1002/ece3.3704

Donaldson, M. R., Burnett, N. J., Braun, D. C., Suski, C. D., Hinch, S. G., Cooke, S. J., & Kerr, J. T. (2016). Taxonomic bias and international biodiversity conservation research. FACETS. https://doi.org/10.1139/facets-2016-0011

Fick, S. E., & Hijmans, R. J. (2017). WorldClim 2: New 1-km spatial resolution climate surfaces for global land areas. International Journal of Climatology, 37(12), 4302–4315. https://doi.org/10.1002/joc.5086

Freiberg, M., Winter, M., Gentile, A., Zizka, A., Muellner-Riehl, A. N., Weigelt, A., & Wirth, C. (2020). LCVP, The Leipzig catalogue of vascular plants, a new taxonomic reference list for all known vascular plants. Scientific Data, 7(1), 416. https://doi.org/10.1038/s41597-020-00702-z

Gal, Y., & Ghahramani, Z. (2016). Dropout as a Bayesian Approximation: Representing Model Uncertainty in Deep Learning. International Conference on Machine Learning, 1050–1059. http://proceedings.mlr.press/v48/gal16.html

Global Biodiversity Information Facility (www.gbif.org). (2019). x(26 August 2019) GBIF Occurrence Download https://doi.org/10.15468/dl.4bijtu.

González-del-Pliego, P., Freckleton, R. P., Edwards, D. P., Koo, M. S., Scheffers, B. R., Pyron, R. A., & Jetz, W. (2019). Phylogenetic and Trait-Based Prediction of Extinction Risk for Data-Deficient Amphibians. Current Biology, 1557–1563. https://doi.org/10.1016/j.cub.2019.04.005

Goodfellow, I., Bengio, Y., & Courville, A. (2016). Deep Learning. MIT Press.

Henry, L., & Wickham, H. (2020). tidyselect: Select from a Set of Strings. https://CRAN.R-project.org/package=tidyselect

Hester, J. (2020). covr: Test Coverage for Packages. https://CRAN.R-project.org/package=covr

Hijmans, R. J. (2018). raster: Geographic data analysis and modeling. https://cran.r-project.org/package=raster

IUCN. (2012). IUCN Red List categories and criteria, version 3.1, second edition. https://portals.iucn.org/library/node/10315

IUCN. (2018). Numbers of threatened species by major groups of organisms (1996–2018). www.iucnredlist.org

IUCN Standards and Petitions Subcommittee. (2017). Guidelines for Using the IUCN Red List—Categories and Criteria. Version 13. Prepared by the Standards and Petitions Subcommittee. Downloadable from http://www.iucnredlist.org/documents/RedListGuidelines.pdf (pp. 1–60).

Kingma, D. P., & Ba, J. (2017). Adam: A Method for Stochastic Optimization. ArXiv:1412.6980 [Cs]. http://arxiv.org/abs/1412.6980

Lang, M. (2017). checkmate: Fast Argument Checks for Defensive R Programming. The R Journal, 9(1), 437–445.

LeCun, Y., Bengio, Y., & Hinton, G. (2015). Deep learning. Nature, 521(7553), 436–444. https://doi.org/10.1038/nature14539

Lughadha, E. N., Bachman, S. P., Leão, T. C. C., Forest, F., Halley, J. M., Moat, J., Acedo, C., Bacon, K. L., Brewer, R. F. A., Gâteblé, G., Gonçalves, S. C., Govaerts, R., Hollingsworth, P. M., Krisai□Greilhuber, I., Lirio, E. J. de, Moore, P. G. P., Negrão, R., Onana, J. M., Rajaovelona, L. R., … Walker, B. E. (2020). Extinction risk and threats to plants and fungi. PLANTS, PEOPLE, PLANET, 2(5), 389–408. https://doi.org/10.1002/ppp3.10146

Lughadha, E. N., Walker, B. E., Canteiro, C., Chadburn, H., Davis, A. P., Hargreaves, S., Lucas, E. J., Schuiteman, A., Williams, E., Bachman, S. P., Baines, D., Barker, A., Budden, A. P., Carretero, J., Clarkson, J. J., Roberts, A., & Rivers, M. C. (2019). The use and misuse of herbarium specimens in evaluating plant extinction risks. Philosophical Transactions of the Royal Society B: Biological Sciences, 374(1763). https://doi.org/10.1098/rstb.2017.0402

Maldonado, C., Molina, C. I., Zizka, A., Persson, C., Taylor, C. M., Albán, J., Chilquillo, E., Rønsted, N., & Antonelli, A. (2015). Estimating species diversity and distribution in the era of Big Data: To what extent can we trust public databases? Global Ecology and Biogeography, 24(8), 973–984. https://doi.org/10.1111/geb.12326

Moat, J. (2017). rCAT: Conservation Assessment Tools. R package version 0.1.5. https://cran.r-project.org/package=rCAT

Monroe, M. J., Butchart, S. H. M., Mooers, A. O., & Bokma, F. (2019). The dynamics underlying avian extinction trajectories forecast a wave of extinctions. Biology Letters, 15(12), 20190633. https://doi.org/10.1098/rsbl.2019.0633

Oliveira, B. F., Sheffers, B. R., & Costa, G. C. (2020). Decoupled erosion of amphibians’ phylogenetic and functional diversity due to extinction. Global Ecology and Biogeography, 29(2), 309–319. https://doi.org/10.1111/geb.13031

Olson, D. M., Dinerstein, E., Wikramanayake, E. D., Burgess, N. D., Powell, G. V. N., Underwood, E. C., D’amico, J. A., Itoua, I., Strand, H. E., Morrison, J. C., Loucks, C. J., Allnutt, T. F., Ricketts, T. H., Kura, Y., Lamoreux, J. F., Wettengel, W. W., Hedao, P., & Kassem, K. R. (2001). Terrestrial ecoregions of the world: A new map of life on Earth. BioScience, 51(11), 933. https://doi.org/10.1641/0006-3568(2001)051[0933:TEOTWA]2.0.CO;2

Ooms, J., & Hester, J. (2020). spelling: Tools for Spell Checking in R. https://CRAN.R-project.org/package=spelling

Parsons, E. C. M. (2016). Why IUCN Should Replace “Data Deficient” Conservation Status with a Precautionary “Assume Threatened” Status—A Cetacean Case Study. Frontiers in Marine Science, 3. https://doi.org/10.3389/fmars.2016.00193

Pebesma, E. J. (2018). Simple features for R: Standardized support for spatial vector data. The R Journal, 10(1), 439–446.

Pelletier, T. A., Carstens, B. C., Tank, D. C., Sullivan, J., & Espíndola, A. (2018). Predicting plant conservation priorities on a global scale. Proceedings of the National Academy of Sciences, 115(51), 13027–13032. https://doi.org/10.1073/pnas.1804098115

Pincheira□Donoso, D., Harvey, L. P., Cotter, S. C., Stark, G., Meiri, S., & Hodgson, D. J. (2021). The global macroecology of brood size in amphibians reveals a predisposition of low-fecundity species to extinction. Global Ecology and Biogeography, n/a(n/a). https://doi.org/10.1111/geb.13287

Polaina, E., Gonzalez-Suarez, M., Kuemmerle, T., Kehoe, L., & Revilla, E. (2018). From tropical shelters to temperate defaunation: The relationship between agricultural transition stage and the distribution of threatened mammals. Global Ecology and Biogeography, 27(6), 647–657. https://doi.org/10.1111/geb.12725

Pollock, L. J., O’Connor, L. M. J., Mokany, K., Rosauer, D. F., Talluto, M. V., & Thuiller, W. (2020). Protecting biodiversity (in all its complexity): New models and methods. Trends in Ecology & Evolution, 35(12), 1119–1128. https://doi.org/10.1016/j.tree.2020.08.015

R Core Team. (2021). R: A Language and environment for statistical computing. R Foundation for Statistical Computing. https://www.r-project.org/

Rapacciuolo, G. (2019). Strengthening the contribution of macroecological models to conservation practice. Global Ecology and Biogeography, 28(1), 54–60. https://doi.org/10.1111/geb.12848

Richards, C., Cooke, R. S. C., & Bates, A. E. (2021). Biological traits of seabirds predict extinction risk and vulnerability to anthropogenic threats. Global Ecology and Biogeography, 30(5), 973–986. https://doi.org/10.1111/geb.13279

Rivers, M. C., Taylor, L., Brummitt, N. A., Meagher, T. R., Roberts, D. L., & Lughadha, E. N. (2011). How many herbarium specimens are needed to detect threatened species? Biological Conservation, 144(10), 2541–2547. https://doi.org/10.1016/j.biocon.2011.07.014

Rolland, J., & Salamin, N. (2016). Niche width impacts vertebrate diversification. Global Ecology and Biogeography, 25(10), 1252–1263. https://doi.org/10.1111/geb.12482

Rondinini, C., Marco, M. D., Visconti, P., Butchart, S. H. M., & Boitani, L. (2014). Update or Outdate: Long-Term Viability of the IUCN Red List. Conservation Letters, 7(2), 126–130. https://doi.org/10.1111/conl.12040

Schmidt, M., Zizka, A., Traoré, S., Ataholo, M., Chatelain, C., Daget, P., Dressler, S., Hahn, K., Kirchmair, I., Krohmer, J., Mbayngone, E., Müller, J. V., Nacoulma, B., Ouédraogo, A., Ouédraogo, O., Sambaré, O., Schuman, K., Wieringa, J. J., Zizka, G., & Thiombiano, A. (2017). Diversity, distribution and preliminary conservation status of the flora of Burkina Faso. Phytotaxa Monographs, 304(1), 1–215.

Silvestro, D., & Andermann, T. (2020). Prior choice affects ability of Bayesian neural networks to identify unknowns. 2005.04987 [Cs, Stat]. http://arxiv.org/abs/2005.04987

Smiley, T. M., Title, P. O., Zelditch, M. L., & Terry, R. C. (2020). Multi-dimensional biodiversity hotspots and the future of taxonomic, ecological and phylogenetic diversity: A case study of North American rodents. Global Ecology and Biogeography, 29(3), 516–533. https://doi.org/10.1111/geb.13050

Stekhoven, D. J., & Bühlmann, P. (2012). MissForest—Non-parametric missing value imputation for mixed-type data. Bioinformatics, 28(1), 112–118. https://doi.org/10.1093/bioinformatics/btr597

Tingley, R., Mahoney, P. J., Durso, A. M., Tallian, A. G., Moran-Ordonez, A., & Beard, K. H. (2016). Threatened and invasive reptiles are not two sides of the same coin. Global Ecology and Biogeography, 25(9), 1050–1060. https://doi.org/10.1111/geb.12462

Ushey, K., Allaire, J. J., & Tang, Y. (2020). reticulate: Interface to “Python.” https://CRAN.R-project.org/package=reticulate

Venter, O., Sanderson, E. W., Magrach, A., Allan, J. R., Beher, J., Jones, K. R., Possingham, H. P., Laurance, W. F., Wood, P., Fekete, B. M., Levy, M. A., & Watson, J. E. M. (2016). Global terrestrial Human Footprint maps for 1993 and 2009. Scientific Data, 3(1), 160067. https://doi.org/10.1038/sdata.2016.67

Walker, B. E., Leão, T. C. C., Bachman, S. P., Bolam, F. C., & Nic Lughadha, E. (2020). Caution needed when predicting species threat status for conservation prioritization on a global scale. Frontiers in Plant Science, 11(April), 1–4. https://doi.org/10.3389/fpls.2020.00520

Walsh, J. C., Venter, O., Watson, J. E. M., Fuller, R. A., Blackburn, T. M., & Possingham, H. P. (2012). Exotic species richness and native species endemism increase the impact of exotic species on islands. Global Ecology and Biogeography, 21(8), 841–850. https://doi.org/10.1111/j.1466-8238.2011.00724.x

Wickham, H. (2011). testthat: Get Started with Testing. The R Journal, 3, 5–10.

Wickham, H. (2020). tidyr: Tidy Messy Data. https://CRAN.R-project.org/package=tidyr

Wickham, H., & Bryan, J. (2020). usethis: Automate Package and Project Setup. https://CRAN.R-project.org/package=usethis

Wickham, H., & Bryan, J. (2021). R Packages (2nd ed.). O’Reilly. https://r-pkgs.org/index.html

Wickham, H., François, R., Henry, L., & Müller, K. (2020). dplyr: A grammar of data manipulation. https://CRAN.R-project.org/package=dplyr

Wickham, H., & Hester, J. (2020). readr: Read rectangular text data. https://CRAN.R-project.org/package=readr

Wickham, H., Hester, J., & Chang, W. (2020). devtools: Tools to Make Developing R Packages Easier. https://CRAN.R-project.org/package=devtools

Xie, Y. (2020). knitr: A general-purpose package for dynamic report generation in R. https://yihui.org/knitr/

Zizka, A., Antonelli, A., & Silvestro, D. (2021). Sampbias, a method for quantifying geographic sampling biases in species distribution data. Ecography, 44(1), 25–32. https://doi.org/10.1111/ecog.05102

Zizka, A., Carvalho, F. A., Calvente, A., Baez-Lizarazo, M. R., Cabral, A., Coelho, J. F. R., Colli-Silva, M., Fantinati, M. R., Fernandes, M. F., Ferreira-Araújo, T., Moreira, F. G. L., Santos, N. M. C., Santos, T. A. B., Santos-Costa, R. C. dos, Serrano, F. C., Silva, A. P. A. da, Soares, A. de S., Souza, P. G. C. de, Tomaz, E. C., … Antonelli, A. (2020). No one-size-fits-all solution to clean GBIF. PeerJ, 8, e9916. https://doi.org/10.7717/peerj.9916

Zizka, A., Silvestro, D., Andermann, T., Azevedo, J., Duarte Ritter, C., Edler, D., Farooq, H., Herdean, A., Ariza, M., Scharn, R., Svantesson, S., Wengström, N., Zizka, V., & Antonelli, A. (2019). CoordinateCleaner: Standardized cleaning of occurrence records from biological collection databases. Methods in Ecology and Evolution, 10(5), 744–751. https://doi.org/10.1111/2041-210X.13152

Zizka, A., Silvestro, D., Vitt, P., & Knight, T. M. (2021). Automated conservation assessment of the orchid family with deep learning. Conservation Biology, 35(3), 897–908. https://doi.org/10.1111/cobi.13616

