## Supplementary tables and figures for "*IUCNN* - deep learning approaches to approximate species’ extinction risk"

Supplementary material S1—Supplementary tables and figures

**Table S1.** Details of custom features, which can be generated by *IUCNN* functions from species’ geographic occurrence records.

| **Feature type** | **Name** | **Feature code** | **Description** |
| --- | --- | --- | --- |
| Geographic | Number of occurrences | tot_occ | The total number of occurrences available for this species |
| Geographic | Number of geographically unique occurrences | uni_occ | The number of geographically unique records available for this species |
| Geographic | Mean latitude | mean_lat | The mean latitude of all records of this species |
| Geographic | Mean longitude | mean_lon | The mean longitude of all records of this species |
| Geographic | Latitudinal range | lat_range | The latitudinal range (.95 quantile - .05 quantile). |
| Geographic | Longitudinal range | lon_range | The longitudinal range (.95 quantile - .05 quantile). |
| Geographic | The hemisphere | alt_hemisphere | 0 = Southern hemisphere, 1 = Northern hemisphere |
| Geographic | Extend of Occurrence | eoo | The extend of occurrence. Calculated by rCAT. For species with less than 3 records set to AOO |
| Geographic | Area of Occupancy | aoo | The area of occupancy, as the sum of area of 4sqkm grid cells, where the species occurs |
| Biome | Tropical & Subtropical Moist Broadleaf Forests | biome_1 | Are at least 5% of the species records present in this biome? |
| Biome | Tropical & Subtropical Dry Broadleaf Forests | biome_2 | Are at least 5% of the species records present in this biome? |
| Biome | Tropical & Subtropical Coniferous Forests | biome_3 | Are at least 5% of the species records present in this biome? |
| Biome | Temperate Broadleaf & Mixed Forests | biome_4 | Are at least 5% of the species records present in this biome? |
| Biome | Temperate Conifer Forests | biome_5 | Are at least 5% of the species records present in this biome? |
| Biome | Boreal Forests/Taiga | biome_6 | Are at least 5% of the species records present in this biome? |
| Biome | Tropical & Subtropical Grasslands, Savannas & Shrublands | biome_7 | Are at least 5% of the species records present in this biome? |
| Biome | Temperate Grasslands, Savannas & Shrublands | biome_8 | Are at least 5% of the species records present in this biome? |
| Biome | Flooded Grasslands & Savannas | biome_9 | Are at least 5% of the species records present in this biome? |
| Biome | Montane Grasslands & Shrublands | biome_10 | Are at least 5% of the species records present in this biome? |
| Biome | Tundra | biome_11 | Are at least 5% of the species records present in this biome? |
| Biome | Mediterranean Forests, Woodlands & Scrub | biome_12 | Are at least 5% of the species records present in this biome? |
| Biome | Deserts & Xeric Shrublands | biome_13 | Are at least 5% of the species records present in this biome? |
| Biome | Mangroves | biome_14 | Are at least 5% of the species records present in this biome? |
| Biome | Lake | biome_98 | Are at least 5% of the species records present in this biome? |
| Biome | Rock and ice | biome_99 | Are at least 5% of the species records present in this biome? |
| Climate | Annual Mean Temperature | bio1 | The median value of this bioclimatic layer for the occurrence records of a species. Records with NA values removed |
| Climate | Temperature Seasonality | bio4 | The median value of this bioclimatic layer for the occurrence records of a species. Records with NA values removed |
| Climate | Mean Temperature of Coldest Quarter | bio11 | The median value of this bioclimatic layer for the occurrence records of a species. Records with NA values removed |
| Climate | Annual Precipitation | bio12 | The median value of this bioclimatic layer for the occurrence records of a species. Records with NA values removed |
| Climate | Precipitation Seasonality | bio15 | The median value of this bioclimatic layer for the occurrence records of a species. Records with NA values removed |
| Climate | Precipitation of Driest Quarter | bio17 | The median value of this bioclimatic layer for the occurrence records of a species. Records with NA values removed |
| Climate | Range of annual Mean Temperature | range_bio1 | The range of value of this bioclimatic layer for the occurrence records of a species. Records with NA values removed. Range is the .95-.05 quantile. |
| Climate | Range of temperature Seasonality | range_bio4 | The range of value of this bioclimatic layer for the occurrence records of a species. Records with NA values removed. Range is the .95-.05 quantile. |
| Climate | Range of mean Temperature of Coldest Quarter | range_bio11 | The range of value of this bioclimatic layer for the occurrence records of a species. Records with NA values removed. Range is the .95-.05 quantile. |
| Climate | Range of annual Precipitation | range_bio12 | The range of value of this bioclimatic layer for the occurrence records of a species. Records with NA values removed. Range is the .95-.05 quantile. |
| Climate | Range of precipitation Seasonality | range_bio15 | The range of value of this bioclimatic layer for the occurrence records of a species. Records with NA values removed. Range is the .95-.05 quantile. |
| Climate | Range of precipitation of Driest Quarter | range_bio17 | The range of value of this bioclimatic layer for the occurrence records of a species. Records with NA values removed. Range is the .95-.05 quantile. |
| Human footprint | Human footprint year 1993 lowest impact | humanfootprint_1993_1 | The fraction of records in areas of the lowest category of human footprint in the year 1993. Footprint was categorized so that categorize represent roughly quantiles. |
| Human footprint | Human footprint year 1993 intermediate impact 1 | humanfootprint_1993_2 | The fraction of records in areas of the second lowest category of human footprint in the year 1993. Footprint was categorized so that categorize represent roughly quantiles. |
| Human footprint | Human footprint year 1993 intermediate impact 2 | humanfootprint_1993_3 | The fraction of records in areas of the second highest category of human footprint in the year 1993. Footprint was categorized so that categorize represent roughly quantiles. |
| Human footprint | Human footprint year 1993 highest impact | humanfootprint_1993_4 | The fraction of records in areas of the highest category of human footprint in the year 1993. Footprint was categorized so that categorize represent roughly quantiles. |
| Human footprint | Human footprint year 2009 lowest impact | humanfootprint_2009_1 | The fraction of records in areas of the lowest category of human footprint in the year 2009. Footprint was categorized so that categorize represent roughly quantiles. |
| Human footprint | Human footprint year 2009 intermediate impact 1 | humanfootprint_2009_2 | The fraction of records in areas of the second lowest category of human footprint in the year 2009. Footprint was categorized so that categorize represent roughly quantiles. |
| Human footprint | Human footprint year 2009 intermediate impact 2 | humanfootprint_2009_3 | The fraction of records in areas of the second highest category of human footprint in the year 2009. Footprint was categorized so that categorize represent roughly quantiles. |
| Human footprint | Human footprint year 2009 highest impact | humanfootprint_2009_4 | The fraction of records in areas of the highest category of human footprint in the year 2009. Footprint was categorized so that categorize represent roughly quantiles. |

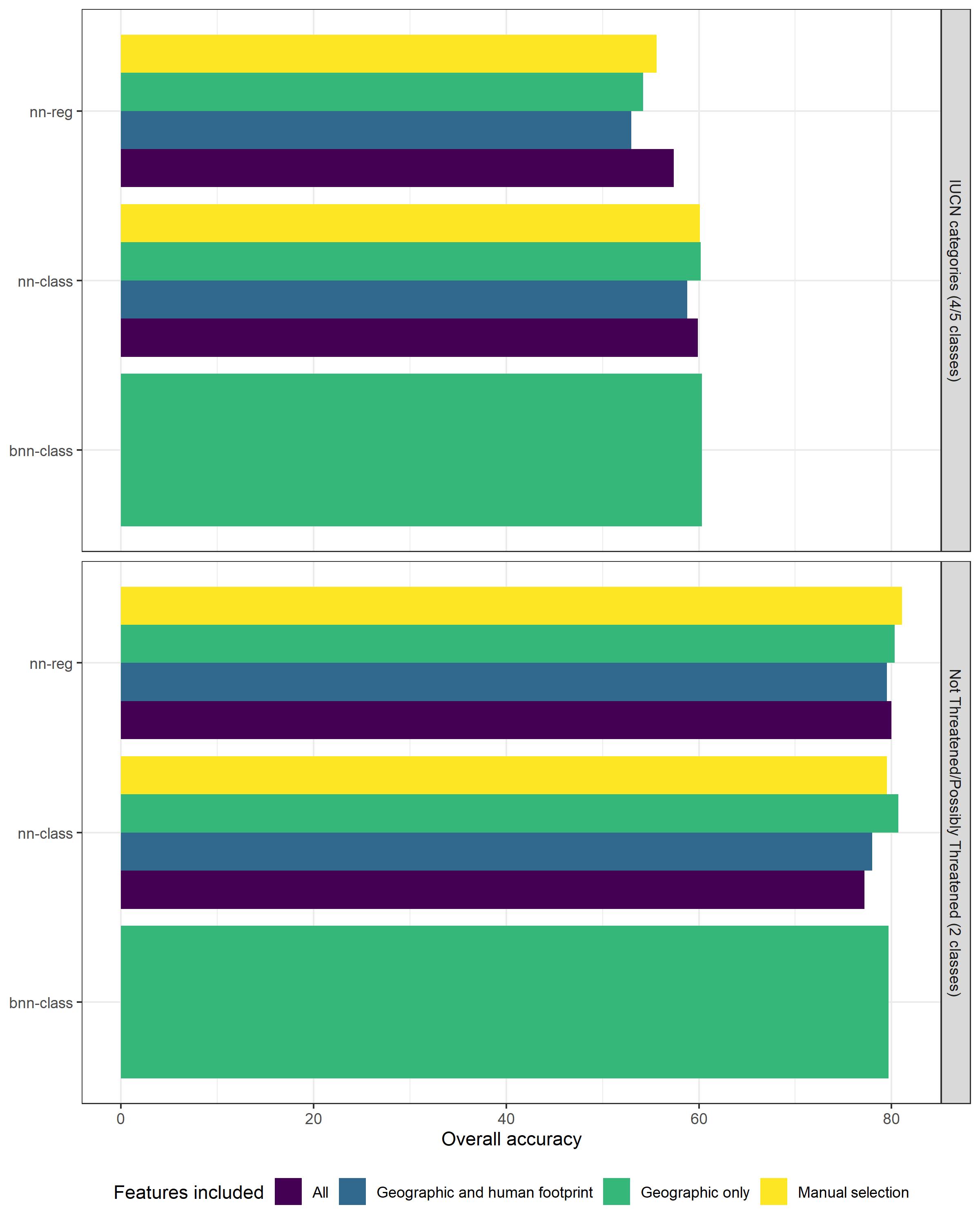

**Figure S1**. The overall accuracy of different neural networks implemented in *IUCNN* in predicting species’ RL status based on a varying set of input features. “All” includes features related to species geographic range (“Geographic”), human footprint at species occurrence (“Human footprint”), species’ climatic niche, and species’ biome occurrence. “Manual selection” includes features from multiple groups selected using the feature importance function. See the vignette distributed with the package for details on the feature groupings.

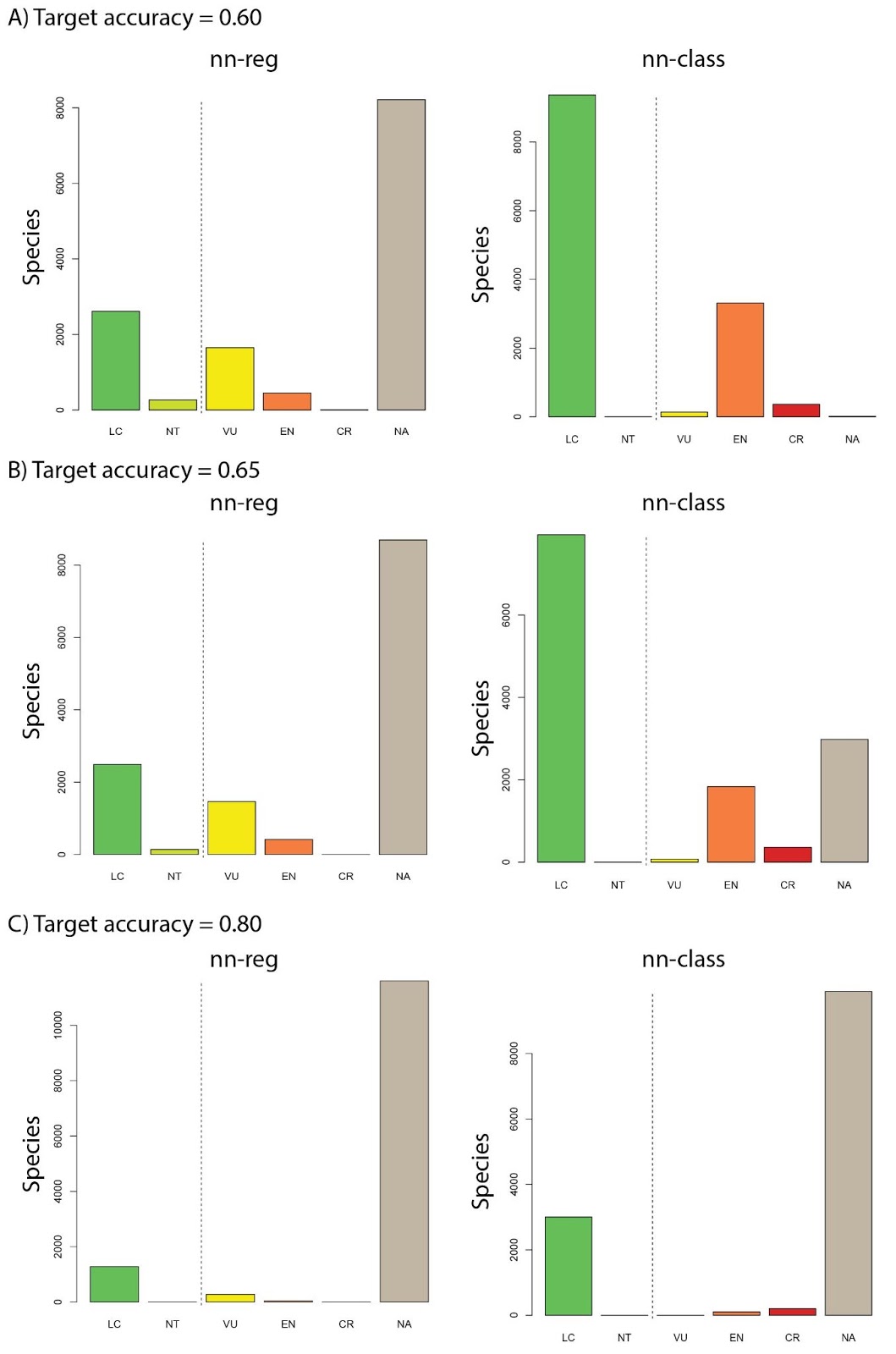

**Figure S2**. Predicted RL categories for 13,207 orchid species with no official assessment available, based on two different *IUCNN* models (left panel: nn-reg, right panel nn-class). **A)** Target threshold of 60% overall accuracy. **B)** Target threshold of 65% overall accuracy. **C)** Target threshold of 80% overall accuracy. The number of assessed species decreases with target accuracy, while even at relatively high accuracy predictive assessments for more than 3,000 species can be made with *nn-class*.

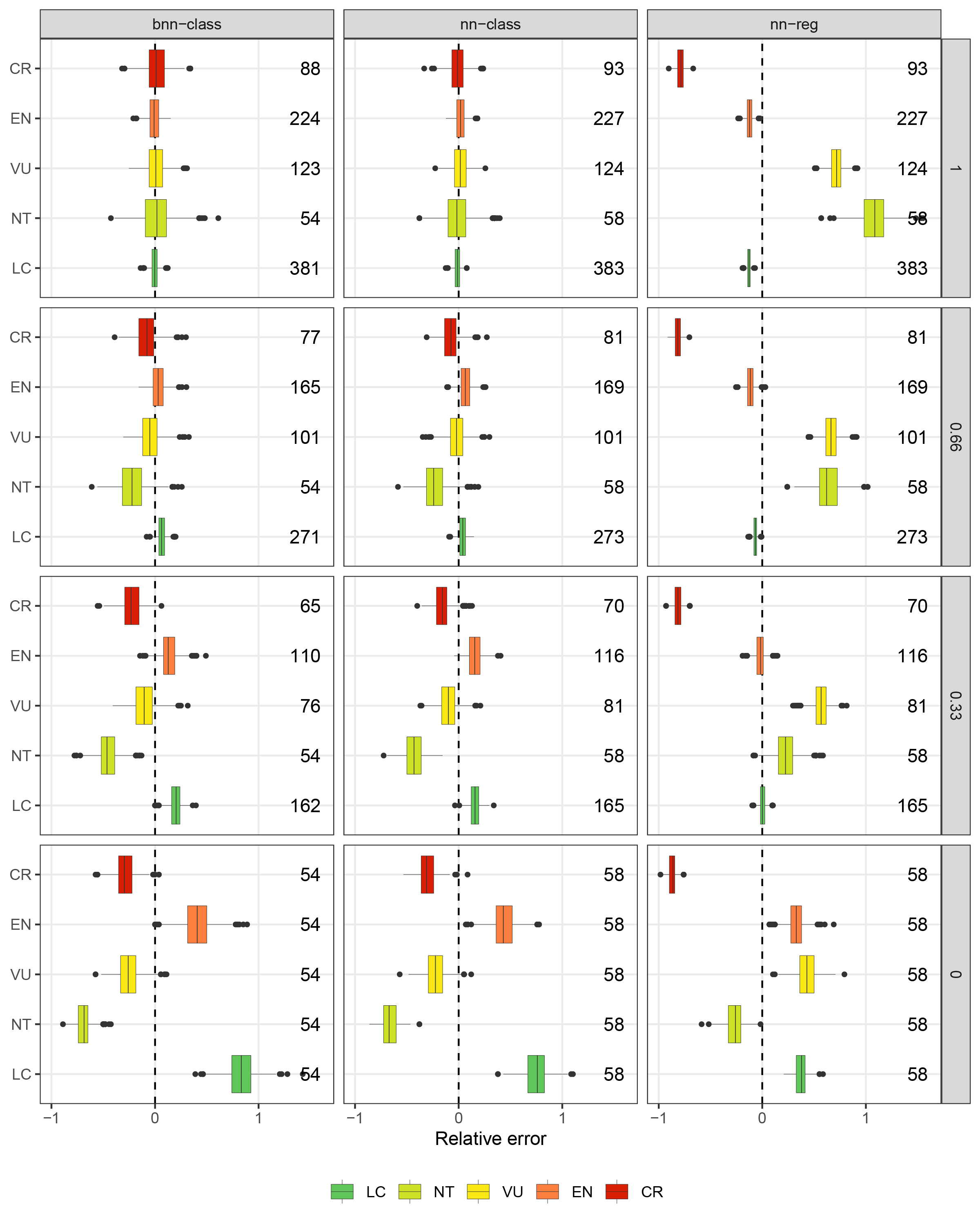

**Figure S3.** The mean absolute error of different *IUCNN* models in predicting the number of species in each extinction risk category depending on class-balance in the training data. We calculated for each test set scenario the relative error of the class distribution predicted by the *IUCNN* model as $RE= \frac{\hat{n}-n}{n}$ where $\hat{n}$ is the number of instances predicted for a given class and $n$ is the true number of instances in that class in the given test set. That is negative values indicate an underrepresentation of a category in the predictions, whereas positive values indicate an overrepresentation. The range of predicted class distributions for each model was produced by repeatedly (n=1000) drawing labels based on the mc-dropout probabilities (posterior probabilities in case of bnn-class) for each instance. We produced increasingly balanced tests sets by applying different balance factors to the test set. A balance factor of 1 leaves the test set unaltered, reflecting the class-distribution in the training data. When set to 0, the instances of all classes are reduced to match those in the minority class (subsampling). In the intermediate steps 0.66 and 0.33, each class is reduced to 66% (33% respectively) of the difference between its original count (balance factor 1) and the minority class count (balance factor 0). That is at bias factor 0, NT, the rarest category, is approximately seven times overrepresented in the test data compared to the training data.
