## Supplementary material for "*IUCNN* - deep learning approaches to approximate species’ extinction risk": IUCNN Tutorial: S2_Approximate_IUCN_Red_List_assessments_with_IUCNN.html


### Approximate IUCN Red List assessments with IUCNN

### Background

The Red List of the International Union for the Conservation of nature (www.iucn.org, IUCN RL), is arguably one of the most thorough and widely used tools to assess the global extinction risk of species. However, the IUCN RL assessment process—usually performed by a group of specialists for each taxonomic group, or professional assessors—are time consuming, and therefore only a small fraction of global biodiversity has been evaluated for the IUCN RL, with a strong bias towards vertebrates and certain regions. These biases and the low fraction of species evaluated, bias conservation towards evaluated groups and prevent synthetic, large-scale ecological and biogeographic analyses of extinction risk.

IUCNN implements neural networks to predict the IUCN status of so far not evaluated or data deficient species based on species traits. IUCNN models are trained on the existing IUCN RL assessments and any traits may be used for prediction, although IUCNN implements a workflow based solely on publicly available geo-referenced species occurrence records and environmental data. Typical examples for the application of IUCNN are to predict the conservation status of a large number of species, to approximate extinction risk or number of threatened species in a region or specific taxonomic group for synthetic analyses or to predict the IUCN category of individual species of interest for systematic or ecological case studies.

```
library(IUCNN)
library(magrittr)
library(dplyr)
```

### Installation

IUCNN uses R and python. All software needed can be installed via R.

1. install IUCNN directly from Github using devtools.

```
install.packages("devtools")
library(devtools)

install_github("azizka/IUCNN")
library(IUCNN)
```

2. Python needs to be installed, for instance using miniconda and reticulated from within R (this will need c. 3 GB disk space). If problems occur at this step, check the excellent documentation of reticulate.

```
install.packages(reticulate)
library("reticulate")
install_miniconda()
```

If python has been installed before, you can specify the python version to sue with `reticulate::use_python()`

3. Install the tensorflow Python module. IUCNN uses functions of the python modules tensorflow and npBNN which also need to be installed (via R).

```
reticulate::conda_install("r-reticulate","tensorflow=2.4")
reticulate::py_install("https://github.com/dsilvestro/npBNN/archive/v0.1.10.tar.gz", pip = TRUE)
```

### Prepare input data

IUCNN predicts the IUCN RL categories of Not Evaluated and Data Deficient species based on geographic occurrence records and a set of training species for which occurrence records and IUCN assessments that are available for a set of reference species (training data). The amount of training species necessary varies with the number of categories but in general “the more, the better”. Ideally, the training dataset should comprise several hundred species or more, so a typical scenario will be to use all available plant species from a region, or all available species from a plant family. If the availability of training species is limited, a good option can be to reduce detail and predict Possibly threatened (IUCN categories “CR”, “EN”, and “VU”) v. Not threatened species (“NT” and “LC”).

Three types of input are necessary, which are easily available for many species:

#### 1. Geographic occurrence records of training species (training occurrences)

Occurrence records might be obtained from a variety of databases, For example, from field collections or public databases such BIEN (https://bien.nceas.ucsb.edu/bien/) or GBIF (www.gbif.org). GBIF data can be obtained from within R via the rgbif package, See here for a tutorial on how to do so. IUCNN needs a dataset with (at least) three columns, containing the species name, decimal longitude coordinates and decimal latitude coordinates. If you are interested in cleaning records from GBIF, you may want to have a look at this blog post and check out the CoordinateCleaner and bRacatus packages.

#### 2. IUCN Global Red List assessment of the training species (training labels)

IUCN RL assessments for the training species can be obtained from IUCN, either via www.iucnredlist.org or via the rredlist package via R (preferred for many species). See here for a tutorial on how to use rredlist. It is important, that all target label classes are well represented in the training data, which is rarely the case for IUCN data, since for instance “VU” and “NT” is rare. If the classes are to imbalanced, consider using possibly threatened (IUCN categories “CR”, “EN”, and “VU”) v. not threatened species (“NT” and “LC”), or the supersampling option of the `train_iucnn` function.

#### 3. Geographic occurrence records of the species for which the IUCN status should be predicted (predict occurrences)

Geographic occurrence for the target species, in the same format as for the training occurrences described above.

Example dataset are available with IUCNN: `data(training_occ)` (training occurrences), `data(training_labels)` (training labels) and `data(prediction_occ)`.

#### Feature preparation

IUCNN uses per species traits to as features for the neural networks. The required input format is a data.frame with one row per species , one column containing the species name and any number of additional columns containing the numerical features for each species. In general, features might represent any trait, for instance from taxonomy (e.g., family), anatomy (e.g., body size), ecology (e.g., feeding guild) or conservation (e.g., population dynamics). However, since often only geographic occurrence data are available IUCNN contains functions to obtain default features from geo-referenced occurrence records alone, by combining them with publicly available environmental data. These default features informing on species range, climatic niche, human footprint and biomes. See Table 2 for a detailed list of all default features. Users may chose to use specific groups of features only via the `type` option of `prep_labels`. In this tutorial, we will use the example datasets from the Orchid family (Orchidaceae) provided with the IUCNN package,

You can prepare the default features with a single call to `prep_features`

```
data("training_occ") #geographic occurrences of species with IUCN assessment
data("prediction_occ")

features_train <- prep_features(training_occ)
features_predict <- prep_features(prediction_occ)
```

#### Label preparation

IUCNN expects the labels for training as numerical categories. So, to use IUCN RL categories, those need to be converted to numeric in the right way. This can be done using the `prepare_labels` function. The function converts the category labels as obtained from the IUCN RL into standardized numeric values, either on the detailed level of IUCN RL categories or the broader Possibly threatened/Not threatened level. See `?prepare_labels` for more information. The labels will be converted into numeric categories following the `accepted_labels` argument, so for instance, in the default case: LC -> 0 and CR -> 4. If you change the accepted labels, the match will change accordingly.

```
data("training_labels")

labels_train <- prep_labels(training_labels)
```

### Running IUCNN

Running IUCNN consists of two steps: 1) training a neural network and 2) predicting the status of new species. IUCNN contains three different neural network approaches to predict the IUCN status of species, which can all be customized. We present the default approach here, see section “Customizing analyses” of this tutorial for details on how to train a Bayesian or regression type neural network.

#### Model training

Based on the training features and labels, IUCNN trains a neural network, via the `train_iucnn` function. There are multiple options to customize the design of the network, including among others the number of layers and the fraction of records used for testing and validation. The `train_iucnn` function will write a folder to the working directory containing the model and return summary statistics including cross-entropy loss and accuracy for the validation set, which can be used to compare the performance of different models.

The following code trains a neural network model with 3 hidden layers of 60, 60, and 20 nodes, with ReLU activation function. By specifying a seed (here, the default: 1234) we make sure the same subsets of data are designated as training, validation and test sets across different runs and model configurations (see below). The model with estimated weights will be saved in the current working directory.

```
res_1 <- train_iucnn(x = features_train,
 lab = labels_train, 
 path_to_output = "iucnn_model_1")
```

The `summary` and `plot` methods give an overview on the training process and model performance.

```
summary(res_1)
#> A model of type nn-class, trained on 702 species and 45 features.
#> 
#> Training accuracy: 0.6
#> Validation accuracy: 0.532
#> Accuracy on unseen data (test set): 0.549
#> 
#> Label detail: 5 Classes (detailed)
#> 
#> Label representation
#>   Label Input_count Estimated_count
#> 1     0          67              92
#> 2     1          12               0
#> 3     2          28               3
#> 4     3          51              80
#> 
#> Confusion matrix (rows test data and columns predicted):
#>    LC NT VU EN CR
#> LC 56  0  1 10  0
#> NT  9  0  0  3  0
#> VU 15  0  1 12  0
#> EN 11  0  1 39  0
#> CR  1  0  0 16  0
plot(res_1)
```

#### Predict IUCN Global Red List status

The trained model can then predict the conservation status of *Not Evaluated* and *Data Deficient* species with the `predict_iucnn` function. The output contains a data frame with species names and numeric labels (as generated with prepare\_labels).

```
predictions <- predict_iucnn(x = features_predict, 
 model = res_1)

plot(predictions)
```

It is important to remember the following when using IUCNN:

1. The resulting IUCNN categories are approximations only. While IUCNN has reached accuracies between 80 and 90% on the broad (threatened v non-threatened) level and up to 80% on the detailed level in some cases, the accuracy may be considerably lower in other cases, which means that some species will be mis-classified.
2. IUCNN is indifferent to the provided features. On the one hand this means that any species traits for which data is available can be used, but on the other hand this means that thought is needed in the choice of the features. The default features of IUCNN are usually a safe choice. The number of features is not limited, but currently IUCNN does not support missing values in the feature table and removes species with missing values.
3. IUCNN is indifferent to the relation between training and test data. So it is possible to use training data from Palearctic birds to predict the conservation status of South American Nematodes. This is not recommended. Instead, a better approach will be to predict the conservation status of species, from training data of the same genus, order, or family. Alternatively, training data could be chosen on geographic region or functional aspects (e.g., feeding guild or body size). However some inclusion of taxonomy/evolutionary history for the choice of training data is recommended.
4. The amount of training data is important. The more the better. Minimum several hundred training species with a more or less equal distribution on the label classes should be included. If training data is limited, the broader Threatened/Not threatened level is recommended.
5. If the proportion of the IUCN RL categories is imbalanced in the training data, the neural networks may be biased towards reproducing these frequencies in the prediction, especially if the imbalance of categories or the difference in category frequencies among training and prediction set are large. To avoid this category frequencies should be balanced in the training data if possible. Otherwise the use of the `supersampling` or option a `nn-reg` type model, or a limitation to the broader Possibly threatened/Not threatened level of detail may remedy the issue.
6. IUCNN predictions are not equivalent to full IUCN Red List assessments. We see the main purpose of IUCNN in 1) identifying species that will likely need conservation action to trigger a full IUCN assessment, and 2) provide large-scale overviews on the extinction risk in a given taxonomic group, for instance in a macro-ecological and macro-evolutionary context.

#### Evaluating feature importance

### Customizing IUCNN analyses

IUCNN contains multiple options to customize the steps of the analyses to adapt the neural networks to the peculiarities of IUCN RL and species distribution data. Below we describe the most important options to customize 1) feature and label preparation, 2) model training and testing, and 3) status prediction. The most important steps and options to customize an IUCNN analysis are summarized in Table 1.

Table 1. Critical steps to customize an IUCNN analysis and relevant considerations at each point.

| Step | Function(s) | Argument | Description | User consideration |
| --- | --- | --- | --- | --- |
| Feature design | prep\_features, feature\_importance | - | Defines the features to be extracted from the provided occurrence records. Available defaults are: biome presence, bioclim variables, human footprint and geographic features. Averaged per species and rescaled. | What determines extinction risk for target group? Which data are available? What do the results of feature importance suggest? |
| Label detail | prep\_labels | level, threatened | Into how many different categories should the species be classified. Can be any number, defaults support full IUCN categories (LC, NT, VU, EN, CR) or binary (Possibly threatened v. Not threatened) | Which detail is needed? Is the accuracy of the detailed level sufficient for the target application? |
| Model type | train\_iucnn | mode | Which model framework should be applied: a categorical classification, a classification taking the ordinal number of categories into account, or a classification based in a Bayesian framework | How many target categories are there? How important is a high accuracy for intermediate classes (e.g. VU, NT, EN)? How important is the uncertainty estimation for each species? |
| Model structure | train\_iucnn | validation fraction | The fraction of the input training data used for validation (v. training). | Which fraction of the data should be used for validation (and hence not training)? How large is the training data? |
| Model structure | train\_iucnn | cv\_fold | The number of folds used for cross-validation. For instance if = 5, the data is divided into 5 folds with 20% of the data used for validation in each run. | How large is the training data? |
| Model structure | train\_iucnn | n\_layers | The number of hidden layers and nodes in the neural network. | How complex should the model be? |
| Model structure | train\_iucnn | balance\_classes | Should the frequency of the class labels in the training data be balanced using supersampling? | How imbalanced are the class labels? Will the frequency of class labels differ between training data and prediction data set? |
| Model structure | train\_iucnn | act\_f\_out/act\_f | The activation function of the neural network | Which relationship between features and labels is expected? |
| Model structure | train\_iucnn | label\_stretch\_factor | The factor to stretch input class labels to | Am I using a nn-reg type model? Does model testing suggest an effect on model accuracy? |
| Model structure | train\_iucnn | drop\_out rate | The number of nodes to be removed in individual epochs. Necessary if a target threshold is to be used with a nn-class model (not bnn-class though) | Is a target accuracy to be sued for prediction? How many nodes are there in the model? |
| Prediction | predict\_iucnn | target\_acc | Defines an overall target accuracy for the model Species which cannot be classified with enough certainty to reach this threshold are labels as data deficient. | Which error rate is acceptable? Which proportion of species needs to be included? |

#### 1) Features and Labels

##### Add and remove feature blocks

The default features are selected based on empirical tests on relevance for different taxa and regions. However, for some analyses only part of the features may be relevant. You can exclude feature blocks using the `type` argument of the `prep_features` function. For instance, to exclude the biome features:

```
features_train2 <- prep_features(training_occ, type = c("geographic", "climate", "humanfootprint"))
```

##### Prepare features individually

If more control over feature preparation is necessary, each feature block can be obtained by an individual function.

Table 2. Functions to obtain default features and options to customize the features.

| Feature block | Function name | Options to customize |
| --- | --- | --- |
| Geographic | `ft_geo` | - |
| Biomes | `ft_biom` | change the reference dataset of biomes (biome\_input, biome.id), remove biomes without any species occurrence (remove\_zeros) |
| Climate | `ft_clim` | the amount of bioclim variables from the default source to be included (type), the resolution of the default input data (res) |
| Human footprint | `ft_foot` | chose the time points from the default source (year), the break points for the different footprint categories (breaks, by default approximately quantiles on the global footprint dataset) or a default source for human footprint (footp\_input) |

For instance:

```
clim_features <- ft_clim(x = training_occ, 
 type = "selected")

clim_features2 <- ft_clim(x = training_occ, 
 type = "all")
```

##### Use custom features

It is also possible to provide features unrelated to the default features. They may contain any continuous or categorical features, but some processing will be needed. The format needs to be a data.frame with a compulsory column containing the species name. Continuous variables should be rescaled to cover a similar range, whereas categorical features should be coded binary (present/absent, as the results of `ft_biom`).

For instance:

```
feat <- data.frame(species = c("Adansonia digitata", "Ceiba pentandra"),
 max_plant_size_m = c(25, 50),
 africa = c(1,1),
 south_america = c(0,1),
 fraction_of_records_in_protected_area = c(25, 75))
```

Table 3. Description of the default features included in `prep_features`. All continuous variables are rescaled to a similar range.

| Feature | Block | Name | Description |
| --- | --- | --- | --- |
| tot\_occ | Geographic | Number of occurrences | The total number of occurrences available for this species |
| uni\_occ | Geographic | Number of geographically unique occurrences | The number of geographically unique records available for this species |
| mean\_lat | Geographic | Mean latitude | The mean latitude of all records of this species |
| mean\_lon | Geographic | Mean longitude | The mean longitude of all records of this species |
| lat\_range | Geographic | Latitudinal range | The latitudinal range (.95 quantile - .05 quantile). |
| lon\_range | Geographic | Longitudinal range | The longitudinal range (.95 quantile - .05 quantile). |
| alt\_hemisphere | Geographic | The hemisphere | 0 = Southern hemisphere, 1 = Northern hemisphere |
| eoo | Geographic | Extend of Occurrence | The extend of occurrence. Calculated by rCAT. For species with less than 3 records set to AOO |
| aoo | Geographic | Area of Occupancy | The area of occupancy, as the sum of area of 4sqkm grid cells, where the species occurs |
| 1 | Biome | Tropical & Subtropical Moist Broadleaf Forests | Are at least 5% of the species records present in this biome? |
| 2 | Biome | Tropical & Subtropical Dry Broadleaf Forests | Are at least 5% of the species records present in this biome? |
| 3 | Biome | Tropical & Subtropical Coniferous Forests | Are at least 5% of the species records present in this biome? |
| 4 | Biome | Temperate Broadleaf & Mixed Forests | Are at least 5% of the species records present in this biome? |
| 5 | Biome | Temperate Conifer Forests | Are at least 5% of the species records present in this biome? |
| 6 | Biome | Boreal Forests/Taiga | Are at least 5% of the species records present in this biome? |
| 7 | Biome | Tropical & Subtropical Grasslands, Savannas & Shrublands | Are at least 5% of the species records present in this biome? |
| 8 | Biome | Temperate Grasslands, Savannas & Shrublands | Are at least 5% of the species records present in this biome? |
| 9 | Biome | Flooded Grasslands & Savannas | Are at least 5% of the species records present in this biome? |
| 10 | Biome | Montane Grasslands & Shrublands | Are at least 5% of the species records present in this biome? |
| 11 | Biome | Tundra | Are at least 5% of the species records present in this biome? |
| 12 | Biome | Mediterranean Forests, Woodlands & Scrub | Are at least 5% of the species records present in this biome? |
| 13 | Biome | Deserts & Xeric Shrublands | Are at least 5% of the species records present in this biome? |
| 14 | Biome | Mangroves | Tropical & Subtropical Moist Broadleaf Forests |
| 98 | Biome | Lake | Are at least 5% of the species records present in this biome? |
| 99 | Biome | Rock and ice | Are at least 5% of the species records present in this biome? |
| bio1 | Climate | Annual Mean Temperature | The median value of this bioclimatic layer for the occurrence records of a species. Records with NA values removed |
| bio4 | Climate | Temperature Seasonality | The median value of this bioclimatic layer for the occurrence records of a species. Records with NA values removed |
| bio11 | Climate | Mean Temperature of Coldest Quarter | The median value of this bioclimatic layer for the occurrence records of a species. Records with NA values removed |
| bio12 | Climate | Annual Precipitation | The median value of this bioclimatic layer for the occurrence records of a species. Records with NA values removed |
| bio15 | Climate | Precipitation Seasonality | The median value of this bioclimatic layer for the occurrence records of a species. Records with NA values removed |
| bio17 | Climate | Precipitation of Driest Quarter | The median value of this bioclimatic layer for the occurrence records of a species. Records with NA values removed |
| range\_bio1 | Climate | Range of annual Mean Temperature | The range of value of this bioclimatic layer for the occurrence records of a species. Records with NA values removed. Range is the .95-.05 quantile. |
| range\_bio4 | Climate | Range of temperature Seasonality | The range of value of this bioclimatic layer for the occurrence records of a species. Records with NA values removed. Range is the .95-.05 quantile. |
| range\_bio11 | Climate | Range of mean Temperature of Coldest Quarter | The range of value of this bioclimatic layer for the occurrence records of a species. Records with NA values removed. Range is the .95-.05 quantile. |
| range\_bio12 | Climate | Range of annual Precipitation | The range of value of this bioclimatic layer for the occurrence records of a species. Records with NA values removed. Range is the .95-.05 quantile. |
| range\_bio15 | Climate | Range of precipitation Seasonality | The range of value of this bioclimatic layer for the occurrence records of a species. Records with NA values removed. Range is the .95-.05 quantile. |
| range\_bio17 | Climate | Range of precipitation of Driest Quarter | The range of value of this bioclimatic layer for the occurrence records of a species. Records with NA values removed. Range is the .95-.05 quantile. |
| humanfootprint\_1993\_1 | Human footprint | Human footprint year 1993 lowest impact | The fraction of records in areas of the lowest category of human footprint in the year 1993. Footprint was categorized so that categorize represent roughly quantiles. |
| humanfootprint\_1993\_2 | Human footprint | Human footprint year 1993 intermediate impact 1 | The fraction of records in areas of the second lowest category of human footprint in the year 1993. Footprint was categorized so that categorize represent roughly quantiles. |
| humanfootprint\_1993\_3 | Human footprint | Human footprint year 1993 intermediate impact 2 | The fraction of records in areas of the second highest category of human footprint in the year 1993. Footprint was categorized so that categorize represent roughly quantiles. |
| humanfootprint\_1993\_4 | Human footprint | Human footprint year 1993 highest impact | The fraction of records in areas of the highest category of human footprint in the year 1993. Footprint was categorized so that categorize represent roughly quantiles. |
| humanfootprint\_2009\_1 | Human footprint | Human footprint year 2009 lowest impact | The fraction of records in areas of the lowest category of human footprint in the year 2009. Footprint was categorized so that categorize represent roughly quantiles. |
| humanfootprint\_2009\_2 | Human footprint | Human footprint year 2009 intermediate impact 1 | The fraction of records in areas of the second lowest category of human footprint in the year 2009. Footprint was categorized so that categorize represent roughly quantiles. |
| humanfootprint\_2009\_3 | Human footprint | Human footprint year 2009 intermediate impact 2 | The fraction of records in areas of the second highest category of human footprint in the year 2009. Footprint was categorized so that categorize represent roughly quantiles. |
| humanfootprint\_2009\_4 | Human footprint | Human footprint year 2009 highest impact | The fraction of records in areas of the highest category of human footprint in the year 2009. Footprint was categorized so that categorize represent roughly quantiles. |

##### Labels: Full categories vs Threatened/Not threatened

The `prep_labels` function may accepted any custom labels as long as they are included in the `accepted_labels` option. It also can provide a classification into threatened/non-threatened, via the `level` and `threatened` options. On the broader level the model accuracy is usually significantly higher.

For instance:

```
labels_train <- prep_labels(training_labels, 
 level = "broad")
```

#### 2) Model training - NN regression model

##### Customizing model parameters

The `train_iucnn` function contains various options to customize the neural network, including among other the fraction of validation and test data, the maximum number of epochs, the number of layers and nodes, the activation function , dropout and randomization of the input data. See `?train_iucnn` for a comprehensive list of options and their description. By default, `train_iucnn` trains a neural network with three hidden layers with 50, 30 and 10 nodes and a sigmoid as activation function. Depending on your dataset different networks may improve performance. For instance, you can set up a different model with 1 hidden layer of 60 nodes, a sigmoid activation function and without using a bias node in the first hidden layer.

```
res_2 <- train_iucnn(x = features_train,
 lab = labels_train, 
 dropout_rate = 0.3,
 path_to_output= "iucnn_model_2",
 n_layers = "60",
 use_bias = FALSE,
 act_f = "sigmoid")
```

You can compare the validation loss of the models using `res_1$validation_loss` and `res_2$validation_loss`. Model 2 in this case yields a lower validation loss and is therefore preferred. Once you chose the preferred model configuration based on validation loss, we can check test accuracy of best model: `res_2$test_accuracy`. The `train_iucnn` function contains various options to adapt the model. See `?train_iucnn` for more detail.

##### Changing the modeling algorithm

There are three neural network algorithms implemented in IUCNN. Besides the default classifier approach based on a tensorflow implementation, these are a Bayesian neural network classifier and a regression type neural network.

The Bayesian approach has the advantage that it returns true probabilities for the classification of species into the relative output classes (e.g. 80% probability of a species to be LC). We consider this approach more suitable for classification of species into IUCN categories, than the default option. It will need more time for model training and should best be applied once you have identified the best model parameters using the default approach. You can run a BNN setting the `mode` option of `train_iucnn` to `"bnn-class"`.

```
res_3 <- train_iucnn(x = features_train,
 lab = labels_train, 
 path_to_output = "iucnn_model_3",
 mode = 'bnn-class')
```

IUCNN also offers the option to train a NN regression model instead of a classifier. Since the IUCN threat statuses constitute a list of ordinal categories sorted by increasing threat level, we can model the task of estimating these categories as a regression problem. Such a model can be trained with the `train_iucnn()` function, specifying to train a regression model by setting `mode = 'nn-reg'`.

```
res_4 <- train_iucnn(x = features_train,
 lab = labels_train, 
 path_to_output = "iucnn_model_4",
 mode = 'nn-reg',
 rescale_features = TRUE)
```

##### Feature importance

The `feature_importance` function can be used to gauge the importance of different feature blocks or individual features for model performance. The function implements the permutation feature importance technique, which randomly reshuffles the values within individual features or blocks of features and evaluate how this randomization affects the models prediction accuracy. If a given feature (or block of features) is important for the models ability to predict, randomizing this feature will lead to a large drop in prediction accuracy. When using `feature_importance` with features other than the default, feature blocks can be defined using the `feature_blocks` option.

```
fi <- feature_importance(x = res_1)
plot(fi)
```

##### Model testing

Before training the final model used for predicting the conservation status of not evaluated species, it is recommended to use the `modeltest_iucnn` function for finding the best settings for your model and dataset. This process, often referred to as hyperparameter tuning, is an essential step for building the most suitable model for the prediction task. The `modeltest_iucnn` function allows you to provide any settings for `train_iucnn` as vectors, which will lead the function to train a separate model for each provided setting. The function will explore all possible permutations of the provided settings, so that the following command results in 9 different models being trained:

```
modeltest_results <- modeltest_iucnn(features,
 labels,
 dropout_rate = c(0.0,0.1,0.3),
 n_layers = c('30','40_20','50_30_10'))
```

The model specifications and settings of each tested model are written to a log-file and can be inspected with the `bestmodel_iucnn` function, to decide which model settings to pick as best model. Different criteria for picking the best model can be selected, such as best prediction accuracy, best predicted over-all status distribution, lowest weighted mis-classification error, etc.

```
best_m <- bestmodel_iucnn(modeltest_results, criterion='val_acc')
```

After model testing, it is necessary to retrain using the model specifications identified by `bestmodel_iucnn`. It is possible to take the respective settings directly from the output of the `bestmodel_iucnn` function, via the `production-model` argument of `train_iucnn`.

```
# Train the best model on all training data for prediction
m_prod <- train_iucnn(train_feat,
                      train_lab,
                      production_model = m_best,
                      overwrite = TRUE)
```

The production model can then be used for the final predictions.

```
# Predict RL categories for target species
pred <- predict_iucnn(pred_feat,
                      m_prod)
plot(pred)
```

#### 3) Status prediction

The `predict_iucnn` function offers options to customize the predictions. The most important option in many cases is `target_acc`, which allows to set an overall target-accuracy threshold that the model needs to achieve. This option is only available for nn-class and nn-reg models that were trained using dropout (see help function of `train_iucnn` for more explanation), as well as for all bnn-class models. The `target_acc` will be achieved by the model being more selective with making a category call for a given instance. All species that cannot be classified with enough certainty to reach this target accuracy will be classified as NA (Not Assessed).

```
pred_2 <- predict_iucnn(x = features_predict, 
 target_acc = 0.7,
 model = res_2)
plot(pred_2)
```

Furthermore, you can turn off the `return_IUCN` option if to return the numerical labels instead of the IUCNN RL category labels.

```
pred_3 <- predict_iucnn(x = features_predict, 
 model = res_2,
 return_IUCN = FALSE)
```

The output of the `predict_iucnn` function is an “iucnn\_predictions” object, that contains several output objects. The predicted labels of the individual instances are accessible with `pred_2$class_predictions` and label probabilities estimated by the neural network via `pred_2$mc_dropout_probs` for “nn-class” and “nn-reg” with dropout, or `pred_2$posterior_probs` for “bnn-class”. For more detail, the `pred_2$raw_predictions` object contains the individual label probabilities resulting from the softmax output layer in case of “nn-class”, or the regressed labels in case of “nn-reg”.

#### 4) The number of species per category

Another stat that can be extracted from the “iucnn\_predictions” object is the overall category distribution predicted for the given prediction instances. This can be accessed with `pred_2$pred_cat_count`, which shows the distribution of the label predictions, including the count of species that could not be predicted given the chosen `target_acc`.

Another stat are the `pred2$sampled_cat_freqs` (only available for dropout models and all “bnn-class” models, see above), which show the class distribution as sampled from the `pred_2$mc_dropout_probs` or the `pred_2$posterior_probs` (for “bnn-class” models). The difference between `pred2$sampled_cat_freqs` and `pred_2$mc_dropout_probs`/`pred_2$posterior_probs` is that the former represents the counts of the best labels determined for each instance, whereas the latter represents labels sampled from the predicted label probabilities, which also proportionally samples the labels for a given instance that do receive the maximum label probability. The latter is done repeatedly to include the stochasticity of the random sampling of classes from the given probability vectors. The `pred_2$mc_dropout_probs`/`pred_2$posterior_probs` can be used to plot histograms of the estimates for each class, and can be reported as uncertainty intervals around the number of species in each class for the set of species that were predicted.
